# Diatoms maintain metabolic activity in deep ocean twilight zones

**DOI:** 10.64898/2026.07.27.741118

**Authors:** Zenghu Zhang, Wenbin Zhao, Shailesh Nair, Da-Zhi Wang, Guangyi Fan, Yongfu Sun, Xue-Wei Xu, Honghai Zhang, Nianzhi Jiao, Thomas Mock, Shanli Mou, Yongyu Zhang

## Abstract

Phytoplankton are primarily confined to the sunlit epipelagic zone, yet intact algal cells persist in the ocean’s mesopelagic twilight zone where light is minimal. However, their survival strategies in the deeper ocean remain unknown. Here, we isolate *Chaetoceros* sp. DS1 from 1000 m depth and reveal its survival mechanisms under prolonged darkness and high pressure, including membrane lipid remodeling and enhanced antioxidant defenses. Importantly, DS1 maintains a latent photosynthetic reserve that activates upon exposure to dim blue light and higher pressure typical of the twilight zone, enabling photosynthesis and nutrient uptake. It exhibits metabolic flexibility by utilizing stored carbon in darkness and assimilating organic carbon under dim light. Global meta-omics data show widespread, transcriptionally active *Chaetoceros*-like diatoms in the twilight zone, expressing blue-light sensors and protein synthesis genes. These findings identify the ocean twilight zone as a previously overlooked niche for metabolically active phytoplankton, expanding understanding of deep-ocean microbial ecology and carbon cycling.

## Introduction

Phytoplankton, the ocean’s primary producers, are predominantly concentrated in the epipelagic zone, where sunlight enables photosynthesis(1). Nevertheless, phytoplankton cells are continuously exported to the deep ocean via cell aggregation, faecal pellet packaging, or downwelling currents(2, 3). This vertical transport of phytoplankton, known as the biological pump, plays a crucial role in oceanic carbon sequestration(4, 5). Traditionally, phytoplankton exported to the deeper ocean including mesopelagic zones (defined as the depth range between 200 to 1000 m) were assumed to arrive only as dead cells or detritus. However, this view is challenged by *in-situ* observations of morphologically intact phytoplankton cells in the dark ocean(6, 7). Intact or viable phytoplankton cells can be detected in bathypelagic marine snow, in aphotic waters below 1000 m, and even in deep-sea sediments(8–13). Hydrostatic pressure did not substantially reduce the abundance of some diatoms, as consuming intracellular stored compounds is one aspect of the energy support for maintaining cellular metabolism (14). These findings raise the possibility that some phytoplankton cells may survive in the deep ocean despite darkness (or having only dim light in twilight zone), low temperatures (0-5°C), and high hydrostatic pressure (HP)(6). Hence, those phytoplankton species would have occupied a niche characterized by reduced microbial degradation and zooplankton grazing(15, 16). A few studies have provided preliminary evidence of distinct phytoplankton taxa occurring exclusively in the twilight zone and even aphotic depths(17, 18), corroborating the hypothesis that at least certain phytoplankton species may be able to survive and possibly be metabolically active in these extreme environments.

Emerging physiological studies provided insight into how they might be adapted to conditions of the deeper ocean. Several traits were identified to facilitate phytoplankton adaptation to the deeper ocean: **Alternative energy acquisition** - facultative heterotrophy(19, 20) enables the utilization of dissolved organic carbon (DOC) as a resource under light limitation. Furthermore, phagotrophy enables some phytoplankton species to benefit from prey such as bacteria as a carbon source(21). **Utilization of residual light** - although photosynthetically active radiation effectively vanishes below ∼200 m, residual blue-green wavelengths can penetrate into the twilight zone. Many phytoplankton species can utilize blue-green light to sustain photosynthetic activity and energy acquisition(22), and Raven *et al.* (23) reported that O_2_-evolving photolithotrophs typically require a minimum photon flux density of ∼10 nmol photons m^-2^ s^-1^ for growth. This supports the idea that residual light (at or below 10^-1^ μmol photons m^-2^ s^-1^) in the twilight zone can support photosynthetic life. Nevertheless, bioluminescent organisms(24) and geothermal emissions of infrared light near hydrothermal vents(25) provide intermittent sources of light for photosynthetic organisms in aphotic zones of the ocean. However, how these light sources contribute to the viability of photosynthetic organisms in the deeper ocean remains elusive. **Dormancy and resting stages** - many diatoms and dinoflagellates can enter metabolically quiescent resting stages that survive extended periods of darkness, low temperature, and nutrient limitation(26, 27).

Despite these proposed mechanisms, the existence of healthy, structurally intact and overall viable microalgal cells in the deeper ocean remains enigmatic. The central barrier to advance our knowledge on a possible deep-sea niche for certain phytoplankton has been the lack of cultivable algal isolates from the deep sea, which would provide experimental evidence and mechanistic insights into adaptive strategies under conditions of the deep sea. To address this limitation, we attempted to isolate phytoplankton strains from the twilight zone (200-1000 m). Unexpectedly, we obtained diverse deep-sea-derived algal strains dominated by diatoms (Fig. S1). One isolated strain, *Chaetoceros* sp. <u>D</u>eep <u>S</u>ea 1 (DS1) was isolated from 1,000 m depth in the Northwest Pacific Ocean and can survive for at least three months under continuous darkness and low temperature, maintaining cellular integrity and viability(28). This cultivable strain provided an unprecedented opportunity to dissect the physiological and molecular basis of diatom persistence in the deep ocean(28–31).

By integrating physiological assays, transcriptomics, and *in-situ* meta-omics, we revealed four main traits that likely underpin DS1’s life in the deep ocean: (1) **tolerance to high pressure**, supported by membrane lipid remodeling and enhanced antioxidant defenses; (2) **a latent “photosynthetic reserve”**, marked by upregulation of photosynthetic complexes and pigments under darkness and high pressure; (3) **activation of photosynthesis when absorbing dim blue light (0.19**`±`**0.02** μ**mol photons m^-2^ s^-1^ blue light)**, enabling nutrient uptake and growth; (4) **facultative mixotrophy**, whereby DS1 relies on stored carbon in darkness and supplements metabolism by taking up dissolved organic carbon when dim blue light becomes available. Beyond laboratory experiments, global oceanic metatranscriptomic data analyses revealed widespread expression of DS1-derived blue-light sensor-cytochrome and rhodopsin genes across the twilight zone, where *Chaetoceros* spp. contribute substantially to eukaryotic ultraphytoplankton communities, at times surpassing epipelagic zones in relative abundances. Thus, our work challenges the traditional paradigm of the twilight zone as a habitat void of active photosynthetic organisms, but is actually a hidden habitat for certain algae with metabolically flexibility.

## Results

### Tolerance of *Chaetoceros* sp. DS1 to prolonged darkness and simulated deep-sea pressure

The deep-sea diatom *Chaetoceros* sp. DS1, isolated from ca. 1000 m depth in the Northwest Pacific, exhibits tolerance to extreme conditions. Previous work has confirmed that this strain can survive three months of darkness and cold stress(28). Consistent with this, we found that after 30 days of darkness, DS1 retained intact vegetative morphology without resting spore formation, and cell abundance remained stable at ∼5 x 10^5^ cells mL^-1^, with no significant decline in cell density (*p* > 0.05; Fig. S1). Notably, DS1 still maintained vegetative integrity even under simulated deep-sea pressure. After 5 days at 5 MPa, 12°C, and darkness (500m-HP-Dark) and 10 MPa, 4°C, and darkness (1000m-HP-Dark), DS1 cells retained unbroken cell morphology and seemingly intact (e.g., homogeneous) cytoplasm (Fig. S2A), with high viability and no resting spore formation (Fig. S3). By contrast, epipelagic strains (*Nannochloropsis* sp. JH01 and *Phaeodactylum tricornutum* CCMP2561) showed a notable decline in photosynthetic efficiency (Fv/Fm) under identical treatment (Fig. S3). Interestingly, similar to DS1, an epipelagic *Chaetoceros* strain (*Chaetoceros muelleri* CCMA318) maintained high viability under simulated deep-sea pressure, indicating that further research is needed to determine how many species of the genus *Chaetoceros* possess the ability to tolerate or adapt to deep-sea environments.

We identified two physiological processes that likely contribute to the viability of DS1 under these deep-sea conditions: First, DS1 increased the total amount of unsaturated fatty acids (UFA) by 24-33%, with specifically increasing C_18:3_ and C_20:5_ by up to 50%, and by producing UFA (C_18:1_, C_16:4_) under these conditions (Fig. 1A). Second, DS1 enhanced the oxidative stress response likely induced by the accumulation of reactive oxygen species (ROS). For instance, the glutathione S-transferase (GST, a key peroxidase) activity under 500m-HP-Dark and 1000m-HP-Dark conditions reached 60 ± 20 U mg^-1^ protein and 46 ± 7 U mg^-1^ protein, a significant increase over surface conditions (26 ± 15 U mg^-1^ protein; *p* < 0.05; Fig. S4).

**Figure 1.**
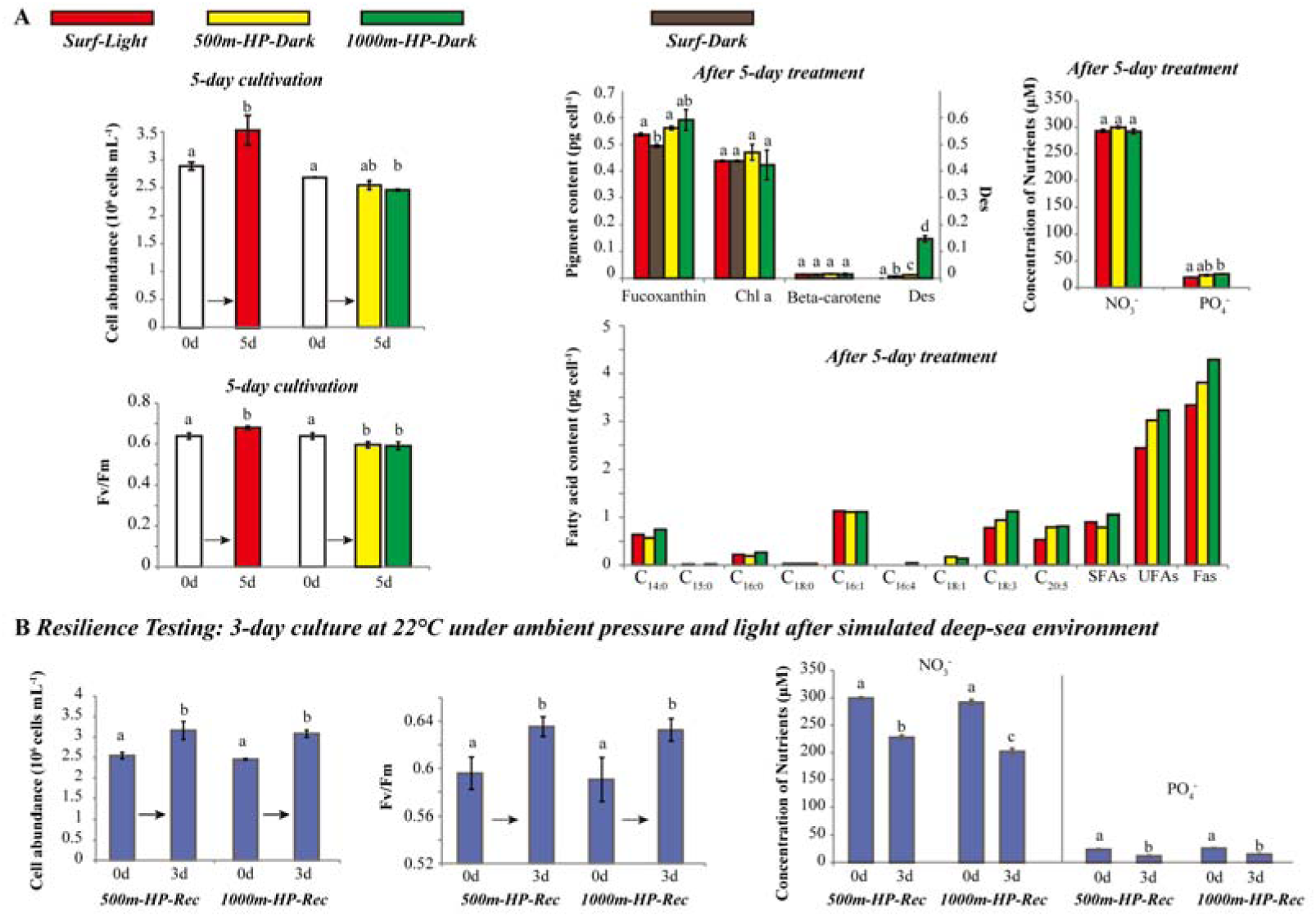
Physiological responses of *Chaetoceros* sp. DS1 under simulated deep-sea pressure and darkness. **(A) Physiological indicators of DS1 after 5-day treatment**: Cell abundance, photosynthetic efficiency (Fv/Fm), pigment content (fucoxanthin, chlorophyll a, β-carotene), de-epoxidation state (Des), inorganic nutrient concentration (NO_3_^-^, PO_4_^3-^), and fatty acid content (SFA: saturated fatty acids, UFA: unsaturated fatty acids). Red column refers to surface control (Surf-Light, white light, 0.1 MPa, 22°C); Yellow refers to treatment mimicking 500 m depth (500m-HP-Dark, darkness, 5 MPa, 12°C); Green refers to treatment mimicking 1000 m depth (1000m-HP-Dark, darkness, 10 MPa, 4°C); Grey refers to dark control (Surf-Dark, darkness, 0.1 MPa, 22°C). **(B) Recovery capacity**: Cell abundance, Fv/Fm, and inorganic nutrient concentration after a 3-day recovery under ambient light/pressure (22°C, 0.1 MPa) following simulated deep-sea exposure. Error bars denote standard deviation; letters indicate statistical significance (Welch’s t-test, *p* < 0.05). For fatty acid content measurements, biological replicates were pooled for a single quantitative analysis due to high biomass requirements, and thus no statistical analysis is presented for this dataset.

After transfer from 500m-HP-Dark and 1000m-HP-Dark conditions to conditions of the surface ocean environment (22°C, atmospheric pressure, light), cell abundance increased significantly (*p* < 0.05) within 3 days, accompanied by a 6-7% increase in Fv/Fm (Fig. 1B). Nutrient uptake resumed, with NO_3_^-^ and PO_4_ ^3-^ concentrations declining significantly (*p* < 0.05; Fig. 1B) over time.

### Survival in darkness and the use of internal carbon reserves

To determine how *Chaetoceros* sp. DS1 survives under prolonged darkness, it was incubated for 30 days under cold and dark conditions (0.1 MPa, 5°C) with or without 100 μM exogenous DOC (Dark-DOC-add vs. Dark-DOC-exc). Cell abundance, photosynthesis parameters (e.g., Fv/Fm), and pigments (chlorophyll *a* and fucoxanthin) exhibited no significant differences between DOC supplementation or its absence (Fig. S2B-C). Thus, supplemented DOC remained unconsumed, with Dark-DOC-add (initial ∼100 μM DOC) maintaining concentrations ∼100-200 μM above Dark-DOC-exc throughout the experiment (Fig. S2D). Instead, DS1 consumed intracellular carbon reserves. Cellular organic carbon decreased from 14.8 ± 1.5 pg cell^-1^ under light to 12.4 ± 0.5 pg cellC¹ after 30 days of darkness (*p* < 0.05; Fig. S2D). The cellular C/N ratio dropped significantly from 8.1 ± 0.3 to 6.3 ± 0.5 in the first 10 days (*p* < 0.05), then stabilized, suggesting catabolism preferentially of carbon over nitrogen reserves (Fig. S2D). This result was corroborated by changes in nutrient concentrations: extracellular phosphate accumulated gradually after day 10 (Fig. S2B), consistent with reduced biosynthetic demand, while dissolved concentrations of extracellular nitrate remained stable (∼50 μM), potentially linked to dissimilatory nitrate reduction to ammonium previously reported for diatoms(32).

Transcriptomic analysis further corroborated the preferential catabolism of internal carbon reserves. Regardless of exogenous organic carbon supplementation, genes responsible for organic carbon uptake (e.g., ABC transporters and solute carriers) exhibited no detectable transcriptional differences (Fig. S5). Under cold and dark conditions, genes related to growth and energy production including glycolysis, photosynthetic antenna proteins, oxidative phosphorylation, nitrogen metabolism, carbon fixation, TCA cycle, and amino acid degradation were downregulated (Table S1, Fig. S6). Upregulation of genes encoding for ribosome biogenesis and UFA biosynthesis suggests that they were required for maintaining viability (Table S2).

### Induction of photosynthetic genes under high pressure and darkness

To further explore DS1’s adaptive mechanisms, we compared its transcriptomic responses under two simulated deeper ocean conditions (500m-HP-Dark and 1000m-HP-Dark) against a surface control (Surf-Light; 0.1 MPa, 22°C, light). An unexpected transcriptional pattern emerged. Although genes related to carbohydrate and lipid metabolism were broadly downregulated, which are consistent with energy conservation under stress, photosynthetic pathways were significantly upregulated under both high pressure and darkness (*p* < 0.05). Upregulated pathways included some of the most essential processes for photosynthetic function, such as genes encoding photosynthetic reaction center proteins (e.g., psbA-C), antenna proteins (e.g., protein fucoxanthin chlorophyll a/c protein), porphyrin/chlorophyll metabolism (e.g., protochlorophyllide reductase), and carotenoid biosynthesis (e.g., phytoene desaturase) (Fig. 2; Table S3). Especially genes encoding key structural and functional components of the photosynthetic apparatus were strongly induced. For instance, *PsbD* (a core subunit of photosystem II, essential for light-dependent reactions) increased 23-fold under 500m-HP-Dark and 8.7-fold under 1000m-HP-Dark (Table S3). Similar trends were observed for PSI subunits, cytochrome b_6_/f complex genes, electron carriers (e.g., *petF*, which encodes ferredoxin), and multiple ATP synthase subunits. Genes involved in pigment biosynthesis were also induced in darkness and high pressure. For instance, carotenoid metabolism genes, including 15-cis-phytoene desaturase (PDS), zeta-carotene desaturase (ZDS), and β-carotene hydroxylase (CrtZ), were upregulated ∼3-fold relative to surface conditions. Photoprotective mechanisms were also induced. For example, the gene encoding diadinoxanthin de-epoxidase (catalyzing the conversion of diadinoxanthin to diatoxanthin in the xanthophyll cycle) was upregulated 4.1-fold at 5 MPa and 7.0-fold at 10 MPa (Table S3).

**Figure 2.**
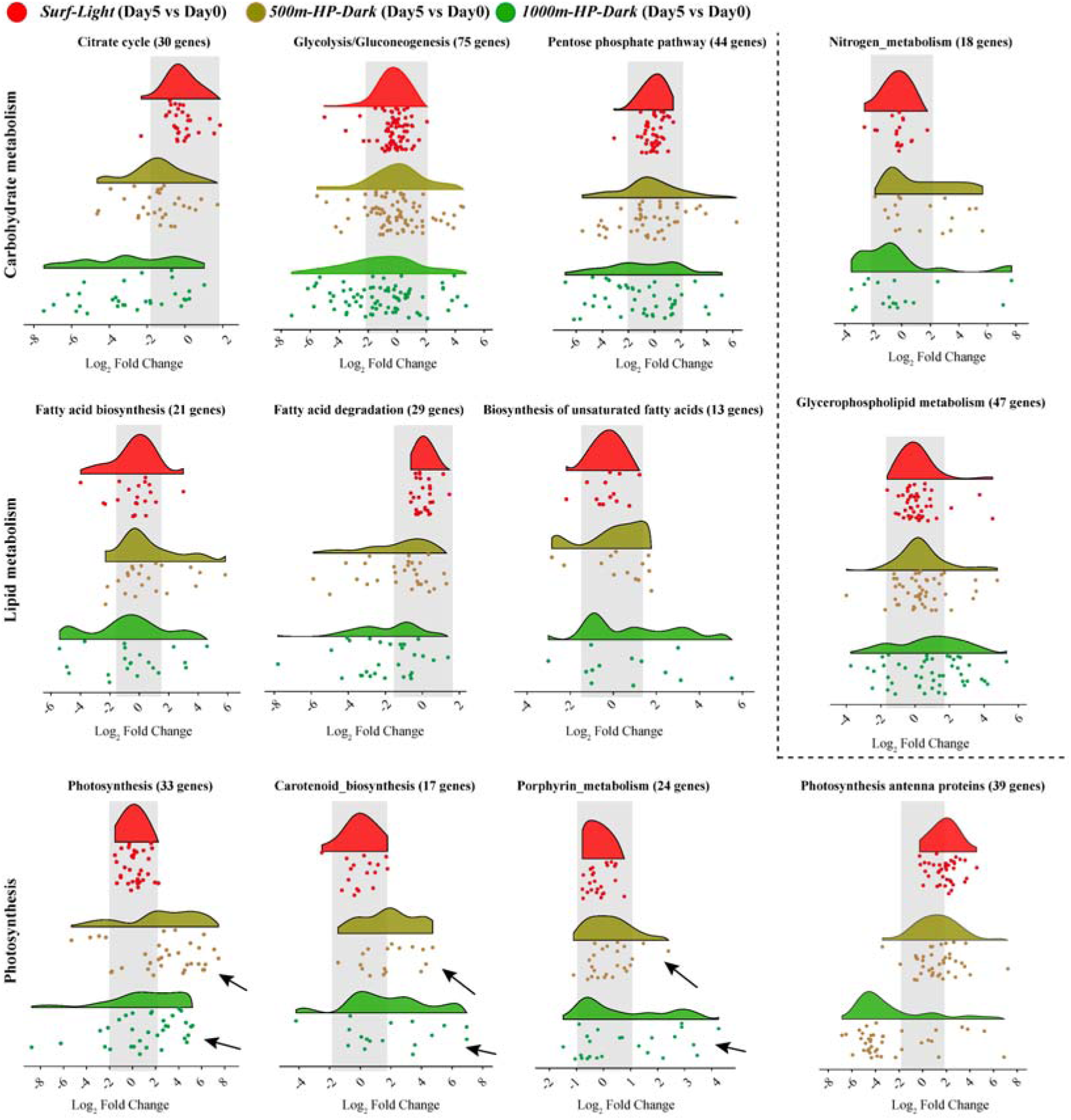
Transcriptomic remodeling of core metabolic pathways in *Chaetoceros* sp. DS1 under simulated deep-sea pressure and darkness. Transcriptomic profiles were compared between Day 5 and Day 0 cultures under different conditions: Red represents surface control (Surf-Light: white light, 0.1 MPa, 22°C), yellow represents treatment mimicking 500 m depth (500m-HP-Dark: darkness, 5 MPa, 12°C), Green represents treatment mimicking 1000 m depth (1000m-HP-Dark: darkness, 10 MPa, 4°C). Each panel depicts gene expression log_2_fold change distributions (violin plots with scatter points) across three functional modules. Carbohydrate and nitrogen metabolism: Citrate cycle, glycolysis/gluconeogenesis, pentose phosphate pathway, and nitrogen metabolism. Lipid metabolism: Fatty acid biosynthesis, fatty acid degradation, biosynthesis of unsaturated fatty acids, and glycerophospholipid metabolism. Photosynthesis and pigment biosynthesis: Photosynthesis (core reaction centers), carotenoid biosynthesis, porphyrin metabolism, and photosynthetic antenna proteins. The gray box represents the gene expression dynamics of DS1 under Surface-Normal conditions. Arrows highlight notable upregulation event under high-pressure and dark conditions compared to surface-normal conditions.

At the physiological level, the level of fucoxanthin significantly increased from 0.49 ± 0.01 pg cell^-1^ under surface pressure and dark conditions to 0.56 ± 0.01 (*p* < 0.05) and 0.59 ± 0.04 pg cell^-1^ under simulated deeper ocean conditions (high pressure, darkness and low temperature) (Fig. 1A). Additionally, the de-epoxidation state (Des) of xanthophyll cycle pigments increased markedly, with a 6-fold rise under 1000m-HP-Dark condition, suggesting enhanced protection of the photosynthetic apparatus against oxidative stress in the extreme environment. Thus, rather than shutting down the photosynthetic machinery in darkness, DS1 maintains and transcriptionally reinforces it, potentially as a pre-emptive mechanism for rapid activation when light becomes available.

### Dim blue light and mixotrophic carbon acquisition in *Chaetoceros* sp. DS1

Residual blue light in the twilight zone ranges from 10^-9^ to 10^-1^ μmol photons m^-2^ s^-1^ (33, 34). DS1’s stronger growth under blue versus white light at ambient pressure (Fig. S7) suggested it may exploit this spectral window to sustain metabolism in deep waters. To test this, we compared two high-pressure treatments, i.e., HP-Blue (5 MPa, 12°C, 0.19±0.02 μmol photons m^-2^ s^-1^ blue light) mimicking twilight-zone light, and HP-Dark (identical pressure and temperature, but no light). This light intensity corresponds to the minimum blue light intensity that can be achieved with the laboratory’s current equipment and conditions. The control groups were set as follows (Fig. S8): white light (denoted as N), darkness (D), and dim blue light (B) under ambient pressure and at 12°C.

#### Transcriptomic signatures of dim blue light-primed mixotrophic carbon acquisition

Dim blue light primes mixotrophic metabolism in DS1 under twilight-zone conditions, as revealed by transcriptomic profiling of core photosynthetic and organic carbon uptake pathways. Relative to HP-Dark, HP-Blue significantly upregulated core photosynthetic pathways, including antenna proteins, carotenoid biosynthesis, porphyrin metabolism, and carbon fixation (*p* < 0.05, Fig. 3B). Key functional genes were strongly induced, including five core photosystem genes critical for light energy capture, 12 chlorophyll a/c-binding protein genes for light harvesting, and *rbcL* encoding Rubisco large subunit (Fig. 3B). Additionally, DS1 cryptochrome genes, encoding specialized blue-light photoreceptors(35), were significantly upregulated under HP-Blue relative to HP-Dark, with a log_2_ fold change of 1.43 (*p* < 0.05, Fig. 4A). Similarly, two rhodopsin-like genes (Rhodopsin-like G protein-coupled receptor and a heliorhodopsin) and were significantly upregulated in the HP-Blue treatment relative to HP-Dark (*p* < 0.05, Fig. 4A). Thus, dim blue light can activate DS1’s latent photosynthetic reserve under twilight-zone conditions. Beyond photosynthesis, HP-Blue induced a broad upregulation of transporters linked to organic carbon uptake. Multiple ABC transporter subfamilies (e.g., ABCA, ABCB, ABCC) and solute carrier transporters for sugars, amino acids, and organic acids were enriched (Fig. 3C).

**Figure 3.**
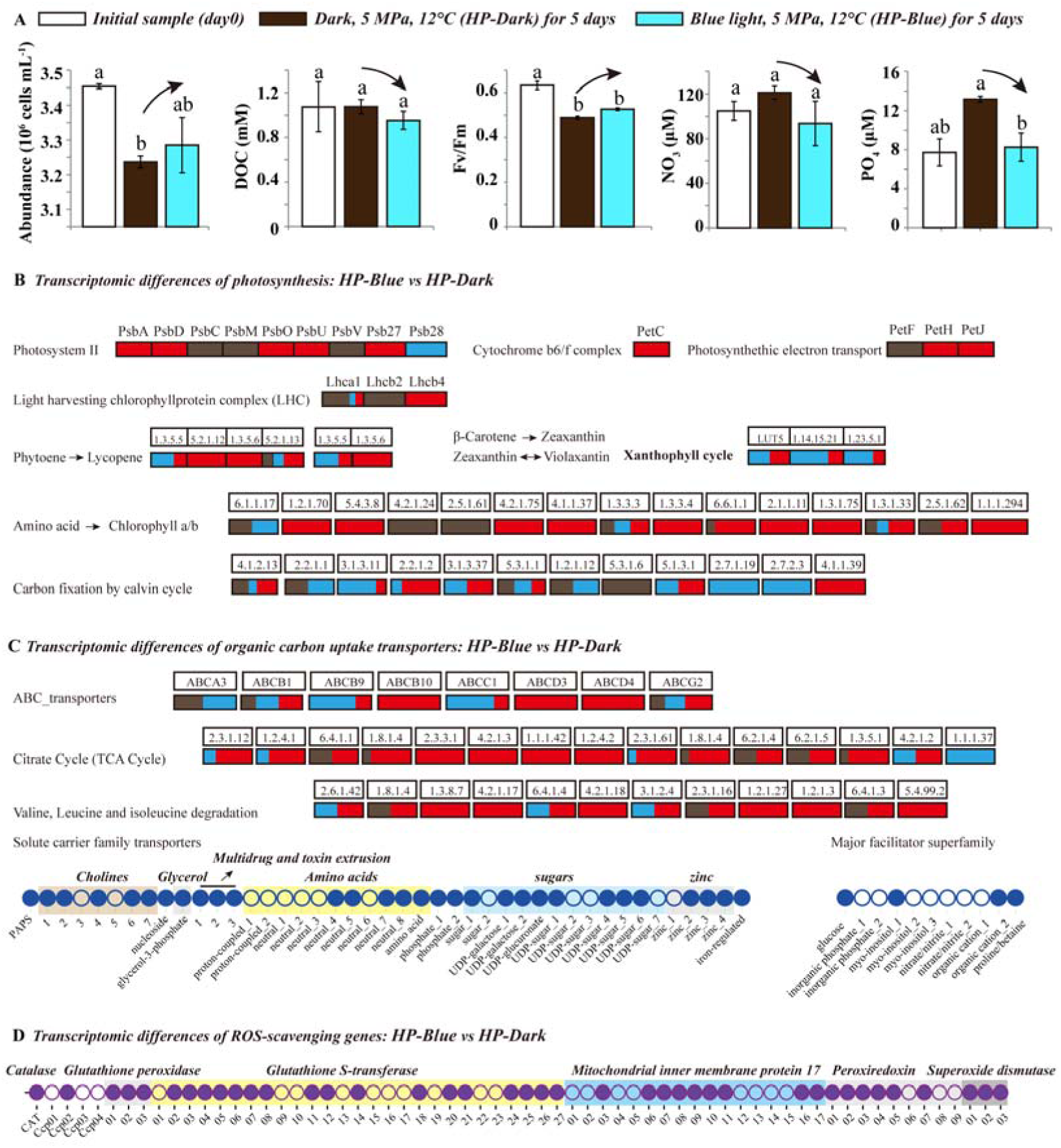
Physiological performance and transcriptional dynamics of *Chaetoceros* sp. DS1 under high-pressure and dim blue light conditions. (A) Physiological responses under simulated 500 m depth conditions (5 MPa, 12°C) in darkness (HP-Dark) or dim blue light (HP-Blue, 0.19 ± 0.02 μmol photons m^-2^ s^-1^). Measured parameters include cell abundance, photosynthetic efficiency (Fv/Fm), dissolved organic carbon (DOC), extracellular inorganic nitrate and phosphate. Letters indicate statistical significance (Welch’s t-test, *p* < 0.05). (B-D) Dim blue light + 5.0 MPa vs darkness + 5.0 MPa. In the boxes, red indicates significant upregulation, blue indicates significant downregulation, and brown indicates non-significant expression change (*p* < 0.05). Each horizontal bar represents a gene cluster, with segment proportions reflecting the fraction of genes in each category. Circles indicate gene-level responses: solid circle indicates significantly upregulated genes; open circle indicates downregulated or non-significantly changed genes (*p* < 0.05). **Functional categories:** (B) Photosynthesis and pigment metabolism: Photosystem I/II, cytochrome b_6_f complex, photosynthetic electron transport, light-harvesting chlorophyll a/b complexes, carotenoid biosynthesis (phytoene to lycopene, xanthophyll cycle), porphyrin metabolism (amino acid to chlorophyll a/b), and carbon fixation. (C) Transporters and central carbon/nitrogen metabolism: ABC transporters, solute carrier family transporters (amino acids, carbohydrates), major facilitator superfamily, tricarboxylic acid cycle, and valine/leucine/isoleucine degradation. (D) Reactive oxygen species scavenging: Glutathione peroxidase (GPx), peroxiredoxin (PRDX), catalase (CAT), cytochrome c peroxidase (CCP), glutathione S-transferase (GST), mitochondrial inner membrane protein 17 (MPV17), and superoxide dismutase (SOD).

**Figure 4.**
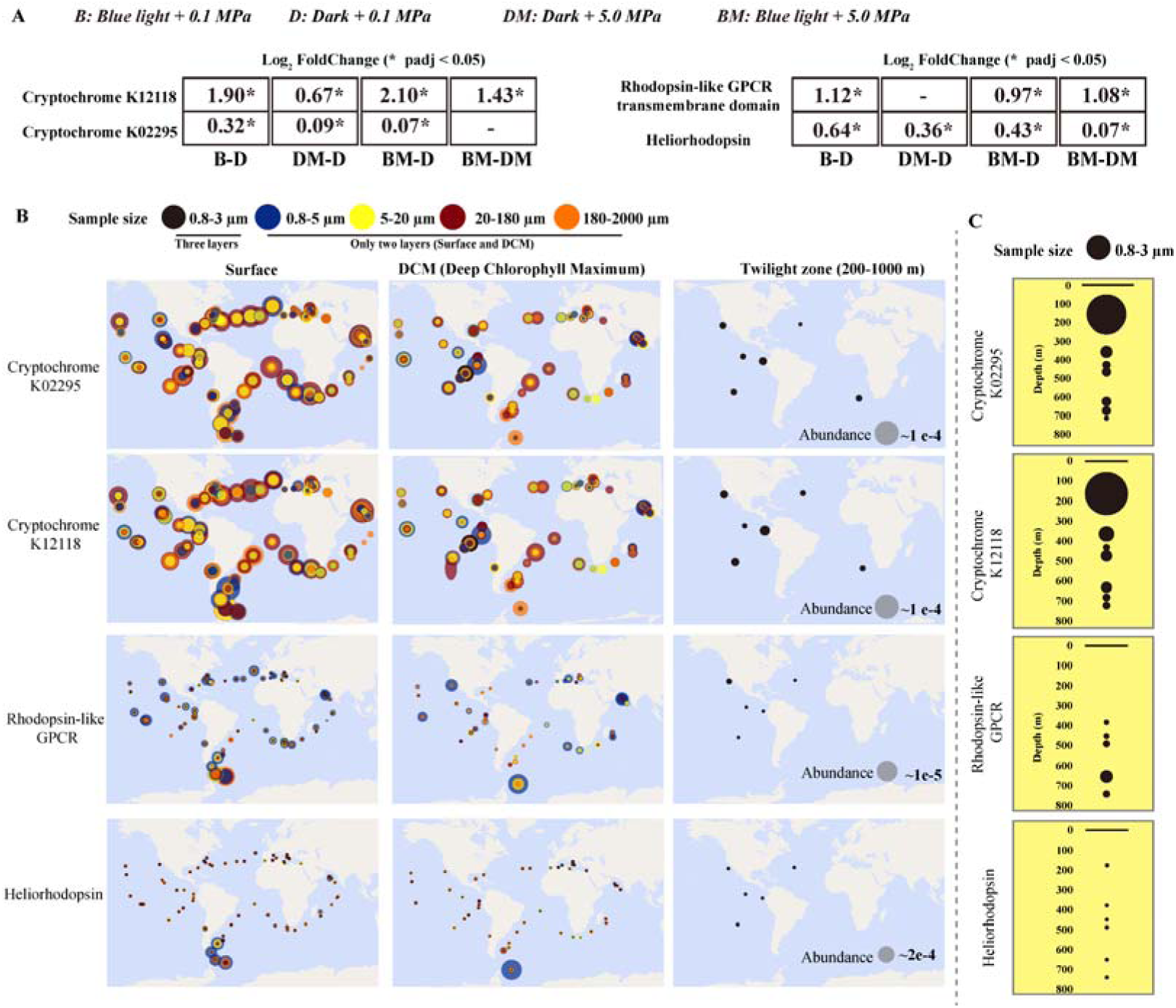
Transcriptional expression of blue-light sensing genes of *Chaetoceros* sp. DS1 in cultures and global oceans. (A) Cryptochrome and rhodopsin gene expression differences (log_2_ fold change) across distinct environments: B-D (dim blue light 0.19 ± 0.02 μmol photons m^-2^ s^-1^ vs darkness, 0.1 MPa), DM-D (darkness, 5 MPa vs darkness, 0.1 MPa), BM-D (dim blue light, 5 MPa vs darkness, 0.1 MPa), BM-DM (dim blue light, 5 MPa vs darkness, 5 MPa). (B) Global distribution of DS-derived cytochrome and rhodopsin transcripts (annotated K02285 and K12118) based on global metatranscriptomes across ocean surface, DCM (Deep Chlorophyll Maximum), and twilight zones (200-1000 m). Circle color and size correspond to transcript abundance (legend denotes sample size and abundance scale). (C) Vertical profiles of DS-derived cytochrome and rhodopsin transcripts in the 0.8-3 μm size fraction, with point size indicating relative abundance.

#### The effect of dim blue light and high pressure on DS1’s stress resilience

Prior studies have documented that deep-sea extreme environment trigger intracellular ROS accumulation and subsequent oxidative damage in marine organisms(36). Transcriptomic results revealed significant upregulation of ROS-scavenging genes, including *GPx* (encoding homologs of glutathione peroxidase), *PRDX* (encoding peroxiredoxin), *GST*, and *MPV17* (encoding mitochondrial inner membrane protein 17) under co-exposure to dim blue light and high pressure (Fig. 3D), indicating that dim blue light activated algal antioxidant defenses under twilight-zone high-pressure environments. Furthermore, given that environmental stress generally triggers dormant states in diatoms(27), we profiled all 91 genes associated with cell cycle progression (Fig. S9). Transcriptomic analysis revealed most of these genes exhibited negligible transcriptional alterations under combined dim blue light and high pressure relative to ambient-pressure darkness or dim blue light alone (|log_2_FoldChange| > 1, *p* < 0.05). Only a small subset showed modest downregulation relative to normal cultures (white light, 0.1 MPa, 22°C) (Fig. S9), demonstrating that twilight-zone condition may not induce DS1 into a dormant state.

#### Physiological validation

Given the extreme experimental conditions, it is challenging to detect physiological changes. Accordingly, no significant changes were observed in key physiological traits under HP-Blue relative to HP-Dark, while a significant change was observed in extracellular inorganic phosphate (PO_4_^3-^) levels (*p* < 0.05, Fig. 3A). Phosphate, as a key nutrient associated with multiple metabolic processes (e.g., ATP synthesis), serves as a critical nutrient for the metabolic activity of DS1 under such conditions. Furthermore, DS1 cell abundance was higher under HP-Blue relative to HP-Dark, accompanied by declines in both DOC and inorganic nutrients (NO_3_^-^, PO_4_^3-^). Fv/Fm values remained stable under high pressure and slightly recovered when the cultures were grown under dim blue light (Fig. 3A), suggesting the functional activation of photosystems. Relative to surface conditions (0.1 MPa, white light or 0.1 MPa, dark), results from the simulated deep-water environment (HP-Blue or HP-Dark) showed that elevated antioxidant enzyme activities (Fig. S4), particularly GST, complemented transcriptional profiles of the underpinning genes and therefore corroborated ROS tolerance.

### Distribution of *Chaetoceros* spp. in the twilight zone and global expression of blue-light receptors

To assess ecological significance of DS1 and its related strains on a global level, we examined global meta-omics datasets covering the surface, deep chlorophyll maximum (DCM), and the twilight zone (see Materials and Methods section)(37). Size-fractionated community analyses targeting ultraphytoplankton (0.2-3 μm) demonstrated that diatoms as ubiquitous and persistent inhabitants of the twilight zone at all 16 sampling stations with paired metagenomic datasets (Fig. 5A). At half of these stations, the relative abundance of diatoms within eukaryotic ultraphytoplankton exceeded their levels detected in surface and DCM layers. *Chaetoceros* dominated the 0.2-3 μm filtered diatom assemblages. Similar to diatom distribution patterns, over half of the stations exhibited higher relative abundances of *Chaetoceros* in the twilight zone than in surface or DCM layers (Fig. 5B, Table S4). Unexpectedly, at two sites, the relative abundance of *Chaetoceros* in the twilight zone reached up to 20% of the eukaryotic ultraphytoplankton (0.2-3 μm cell diameter). At one North Atlantic station (Stations Tara_205), the relative abundance of *Chaetoceros* in the twilight zone reached ∼20%, while its surface abundance remained at 15%. Similarly, at least one station in the mesopelagic zone of the Southern Ocean (Stations Tara_065) exhibited a relative abundance of *Chaetoceros* spp. of over 20%, with its surface abundance remaining below 1%. Consistent with this global trend, our *in-situ* 18S rRNA transcript sequencing across vertical profiles at two open-ocean stations (Stations D1 and D6) in the Northwestern Pacific Ocean confirmed the active presence of *Chaetoceros* spp. in the twilight zone. At Station D1, relative abundance of *Chaetoceros* and other diatom genera peaked at 250 m (a typical upper twilight zone depth with faint blue light penetration), with a substantial increase observed below the DCM layer (85 m) and they were sustained at depths as deep as 800 m. Similarly, at Station D6, *Chaetoceros* spp. exhibited its highest relative abundance in the twilight zone at 1000 m, and was frequently detected at depths exceeding 250 m, where environmental light was below the detection limit (Fig. S10).

**Figure 5.**
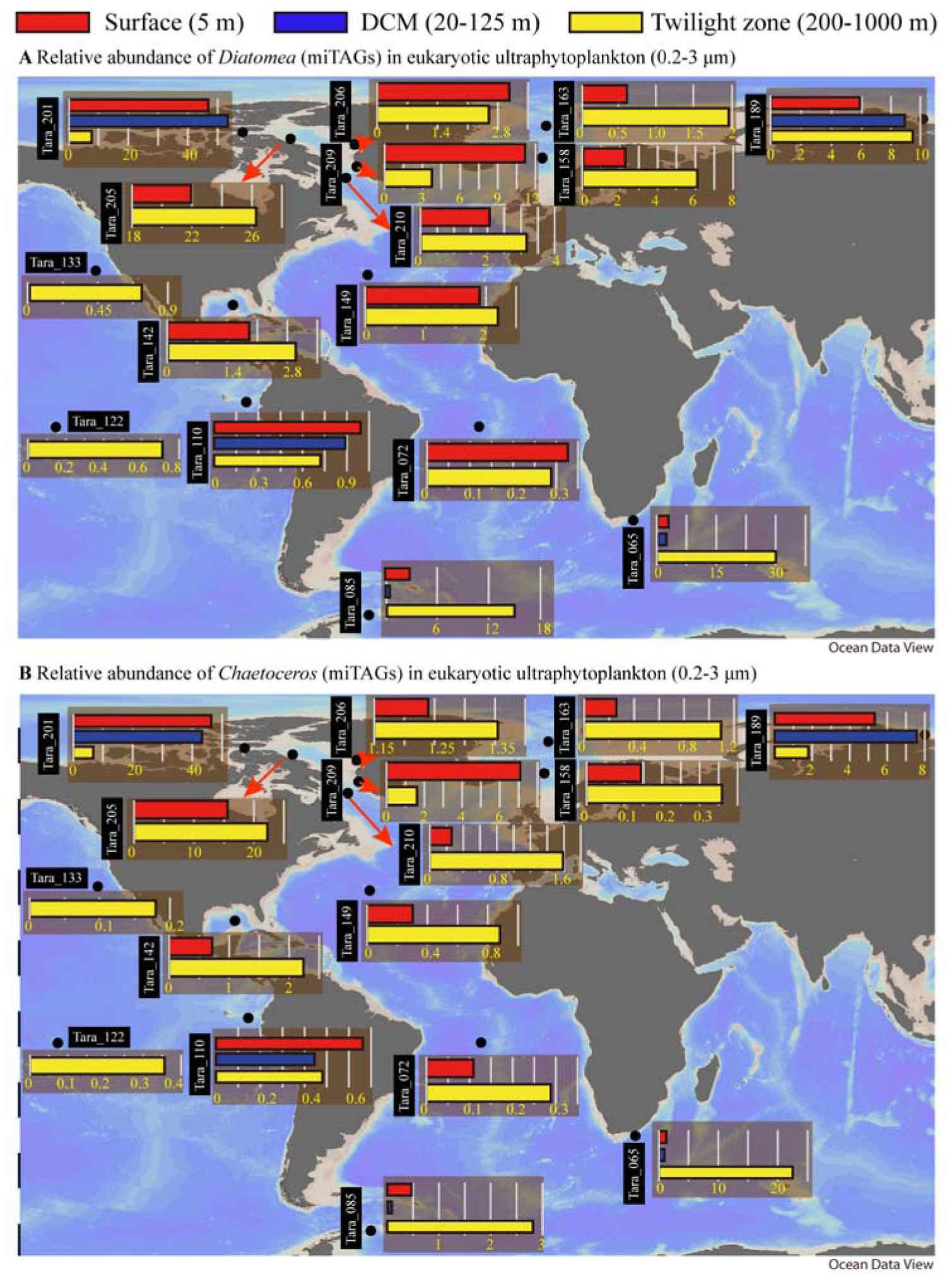
Global and vertical distribution of *Diatomea* (A) and *Chaetoceros* (B) in the ocean. Color legend: red refers to the surface layer (5 m), blue refers to the Deep Chlorophyll Maximum (DCM, 20-125 m), and yellow refers to the twilight zone (200-1000 m). The map illustrated the relative abundance of *Diatomea* and *Chaetoceros* miTAGs (Tara Oceans metagenomic data for species-level taxonomic profiling) within eukaryotic ultraphytoplankton (0.2-3 μm) communities across global ocean sampling sites.

To further verify the active survival of *Chaetoceros* in the twilight zone, we performed metatranscriptomic analysis of the Tara Oceans dataset using PF00009 (i.e., GTP-binding elongation factor family, EF-Tu/EF-1A subfamily) as a marker. This Pfam family is among the most highly expressed diatom-associated gene families in the global ocean(38). EF-1A is ubiquitously present in eukaryotes, where it plays a pivotal role in mediating protein synthesis(39), and it has been marker for phylogenetic studies(40). Studies have reported that the expression of EF-1A in plant cells is activated upon entry into the cell cycle (during the transition from quiescent cells to dividing cells) and continues during the transit through the cycle(41), which suggests it as a marker to characterize metabolic activity. Our analysis revealed widespread expression of PF00009 throughout the water column, including the twilight zone (Fig. 6A and 6B), with *Stramenopiles* (including diatoms) as the dominant group. Within *Stramenopiles*, diatoms exhibited relatively high levels in the twilight zone. Unexpectedly, *Chaetoceros* was notably present in the twilight zone at all but one station, indicative of transcriptional activity in this twilight zone. We further observed that *Chaetoceros* showed increased relative abundance in the twilight zone at all but one station, while exhibiting extremely low relative abundance in the surface and DCM layers at numerous sites (Fig. 6C-E). However, as samples from the twilight zone were collected from the 0.8-3 μm size fraction, which differs from the size fraction (0.8-5 μm) used for surface and DCM layers, this finding warrants further validation. Similarly, homologs of two DS1-derived blue-light photoreceptor (cryptochrome and rhodopsin) genes in Tara Oceans metatranscriptomic datasets(42) showed widespread expression in twilight-zones (Fig. 4B).

**Figure 6.**
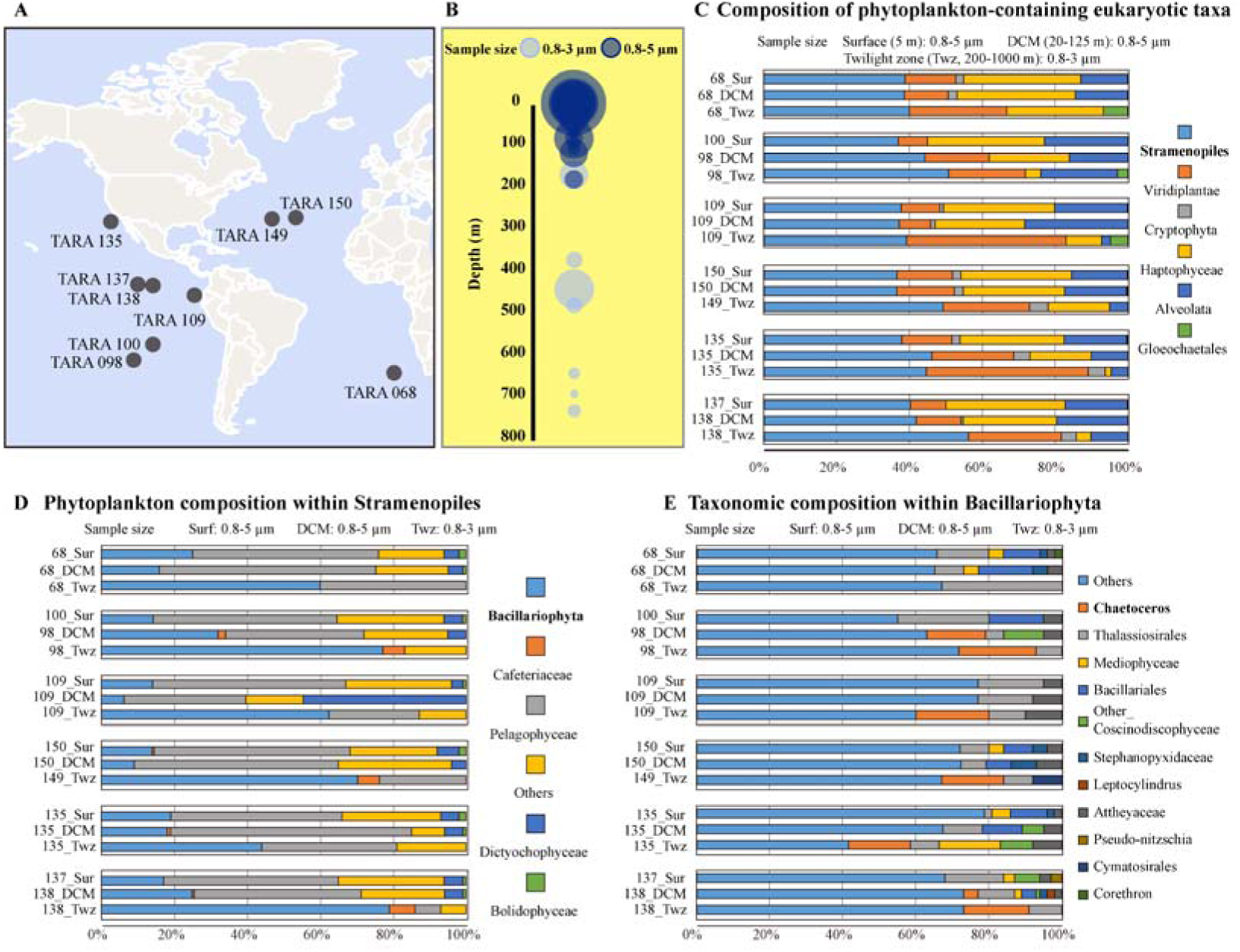
Eukaryotic phytoplankton (0.8-3/5 μm) community structure inferred from global ocean metatranscriptomic data using Pfam family-PF00009 (GTP-binding elongation factor family, EF-Tu/EF-1A subfamily) searches. (A) Geographical distribution of Tara Oceans sampling stations analyzed in this study. (B) Vertical distribution of PF00009-annotated reads across different water layers; bubble size represents the relative abundance (percent of total reads) of PF00009-annotated reads, and color indicates distinct particle size fractions. (C) Relative abundance of major eukaryotic phytoplankton taxa across sampling stations and water layers (Surface, Chlorophyll Maximum layer, and Twilight zone), identified via PF00009 annotation. (D) Phytoplankton community composition within *Stramenopiles* (PF00009-based taxonomic classification). (E) Taxonomic composition within *Bacillariophyta* (PF00009-based taxonomic classification). Sample size fraction: 0.8-5 μm for surface and DCM layers, and 0.8-3 μm for the twilight zone. Selection of different size-fractionated samples was due to the 0.8-5 μm fraction not being sampled in the twilight zone, whereas no samples or annotated data for the 0.8-3 μm fraction were available for the surface and DCM layers. Where data for specific layers at certain stations were missing, they were supplemented with data from adjacent stations. Abbreviations: Surf, Surface seawater; DCM, Deep Chlorophyll Maximum layer; Twz, Twilight zone.

## Discussions

Historically, the presence of phytoplankton in the deeper ocean (> 200 m) including the twilight zone has been attributed to sinking but not to a niche for species adapted to higher pressure and dim blue light, although microscopy and molecular surveys(7, 17) provided first evidence. However, this hypothesis was difficult to test because there was no cultivable deep-sea phytoplankton strain available. *Chaetoceros* sp. DS1, a diatom isolated from 1000 m depth in the Northwest Pacific fills this gap and provides first insights into mechanisms underpinning its adaptation to the twilight zone. The main limitations in the twilight zone for photosynthetic organisms are dim blue light, increased pressure, and overall low temperatures. Our results have shown that DS1 and possibly other *Chaetoceros* species such as the tested *C. muelleri* from the epipelagic ocean appear to have an adaptive response to these conditions (Fig. S3). Thus, it appears that the ability to survive as a photosynthetic diatom in the twilight zone has evolved in the genus *Chaetoceros*, but more species from different diatom genera and the natural environment need to be tested to corroborate this hypothesis.

DS1 instead not only showed resilience under higher pressure and dim blue light characteristic of twilight-zone conditions but it even activated some genes involved in photosynthesis even under complete darkness, maintained photosynthetic pigments, membrane integrity and enhanced antioxidant defenses (Figs. 2, S3 and S4). In addition, DS1 appears to catabolize internal carbon stores (Fig. S2D) and may utilize external DOC if sufficient light is available for active transport and metabolism, based on the significant upregulation of cryptochrome genes, rhodopsin genes, transporters and solute carriers (*p* < 0.05), coupled with the observed downward trend in DOC concentrations (Figs. 3B and 4A). The latter appears to be the case at already 0.19 ± 0.02 μmol photons m^-2^ s^-1^ blue light. Although, catabolism of internal carbon stores appears to be common for diatoms under energy limitations such as during the polar night or the formation of resting stages, to be physiological active under 0.19 ± 0.02 μmol photons m^-2^ s^-1^ blue light has so far been observed for polar microalgae(43), which live under similar environmental conditions except elevated pressure, this key difference in pressure exposure distinguishes the adaptive strategy of the deep-sea diatom. Unlike polar diatoms, which only need to retain intact photosynthetic machinery to make use of low intensity blue light, DS1 integrates low-light adaptation with pressure resilience. DS1 remodels its cell membranes by increasing unsaturated fatty acids to prevent pressure-induced rigidity (Fig. 1A), enhances antioxidant defenses to reduce pressure-related reactive oxygen species (Fig. S4), and even upregulates genes for photosynthetic complexes and pigment biosynthesis when in darkness and under high pressure (Fig. 2), maintaining a “photosynthetic reserve” that activates quickly once dim blue light becomes available.

As a critical model for the adaptation of photosynthetic organisms in the twilight zone, our data show that DS1 survives in the twilight zone via multiple interconnected traits including tolerance to pressure and darkness, a latent “photosynthetic reserve” that is transcriptionally primed under stress, and dim blue-light-driven mixotrophy. Combined with the global distribution of *Chaetoceros* spp. and active expression of DS1-derived genes in the twilight zone, these features highlight the ecological relevance of DS1-like adaptations and therefore make a case for reassessing the depth limits of active phytoplankton and therefore likely also carbon cycling in the ocean’s interior.

### Non-canonical photosynthetic activation under high pressure and darkness

Conventional paradigms hold that high hydrostatic pressure, low temperature, and darkness suppress algal photosynthesis in the deep ocean(44). Yet, DS1 shows a non-canonical response. High pressure under dark conditions likely serves as signals for DS1 to adapt and rapidly reactivate metabolism when light reappears through hydrodynamic processes or residual transmission of photons in the blue spectrum. In addition to this photosynthetic activation, high pressure also induced membrane lipid remodeling via enhanced unsaturated fatty acid biosynthesis (Fig. 1A), elevated oxidative stress responses, as evidenced by the significant induction of glutathione S-transferase activity (*p* < 0.05, Fig. S3) to maintain cellular integrity and metabolic readiness. These strategies are particularly suited to the twilight zone where blue photons may penetrate and therefore serve as a light source for photosynthesis.

### Exploiting dim blue light in the twilight zone

Blue light penetrates deepest into the ocean and deep-sea bacteria have been shown to sense and utilize this light source to support growth(45, 46). DS1’s dominant pigment, fucoxanthin, efficiently absorbs in the 440-540 nm range, aligning with the spectral profile of light in the twilight zone(47, 48). Based on the results of our experiments with a light intensity of 0.19 μmol photons m^-2^ s^-1^, which falls within the twilight zone range ≤ 10^-1^ μmol photons m^-2^ s^-1^, DS1 should take advantage of this residual light in the twilight zone. Compared to the darkness condition (D), the two cryptochrome genes (K12118 and K02295), common blue light sensors(35), both are upregulated under high pressure in darkness (DM), and under the combined dim blue light and high pressure (BM) (Fig. 4A). Minor discrepancies in their expression between high pressure under dim blue light and high pressure in darkness (BM-DM) may arise from complex interactions between these two stimuli. Importantly, both cryptochromes were expressed under the simulated deep-sea conditions, consistent with our global meta-omic data showing cryptochrome transcription in twilight zones (Fig. 4B), supporting their role in environmental adaptation. Here, the increased expression of cryptochromes suggests that blue photons may trigger extensive signaling cascades underpinning adaptive responses. Furthermore, by comparing the high-pressure dark (HP-Dark) and high-pressure blue light (HP-Blue) treatments (Figs. 3 and 4), blue light activated the latent photosynthetic machinery primed by pressure and mixotrophic metabolism. Twilight-zone condition (HP-Blue) significantly stimulated the antioxidant defenses compared to Dark treatments (*p* < 0.05, Fig. S4). Reinforced by antioxidant defenses (e.g., glutathione metabolism, xanthophyll cycling), which appear to be characteristic for many deep-sea organisms(36), DS1 and its relatives represent a group of phytoplankton capable of exploiting the twilight zone niche, an environment previously considered insignificant for photosynthetic organisms.

In considering the ecological relevance of light exposure for DS1 cells in the twilight zone, the 0.19 μmol photons m^-2^ s^-1^ dim blue light used acts as a practical experimental irradiance to characterize DS1’s photosynthetic activation. This value does not represent the minimum light threshold for a functional response and DS1 is likely capable of light-dependent activation at even lower photon fluxes in natural twilight zone waters. Natural irradiance in the twilight zone ranges from 10^-9^ to 10^-1^ μmol photons m^-2^ s^-1^(49), and the universal minimum light threshold for photosynthetic carbon fixation in phototrophs is estimated to be 0.01 μmol photons m^-2^ s^-1^ (23), a flux far below our experimental setting. Light penetration to twilight zone depths is primarily regulated by water column transparency: in high-transparency pelagic waters (attenuation coefficient K=0.025 m^-1^), an irradiance of 10^-1^ μmol photons m^-2^ s^-1^ (near our experimental value) can reach 370 m, a typical upper twilight zone depth, while even in low-transparency coastal waters (K=0.06 m^-1^), this photon flux penetrates to 150 m, the upper boundary of the twilight zone(49). Vertical mixing, eddy diffusion, and upward advection may transport DS1 cells to the upper twilight zone (200-300 m) or the euphotic zone(34), where light exposure naturally approaches or reaches our experimental irradiance of 0.19 μmol photons m^-2^ s^-1^. Notably, DS1 encodes rhodopsin genes that are activated under condition mimicking 500 m depth and dim blue light (Fig. 4). Rhodopsins if they contribute to the synthesis of ATP through proton pumping therefore might contribute to energy production under low-light conditions, further supporting the probability that DS1 can respond to photon fluxes lower than our experimental irradiance.

### Conditional DOC utilization

Uptake of DOC is considered to be a survival mechanism for deep-sea phytoplankton(18). However, DS1 employs a more nuanced strategy in terms of exploiting DOC from the deep sea. During prolonged darkness, exogenous DOC did not have any measurable effect on its activity. Instead, DS1 sustains viability by catabolizing internal carbon reserves, similar to polar diatoms enduring long dark winters(31, 50). Strikingly, the presence of dim blue light triggered a metabolic shift in DS1. ABC transporters and solute carriers were upregulated (Fig. 3C), supporting the assimilation of exogenous DOC under conditions in the twilight zone. This dim blue-light dependent mixotrophy echoes strategies seen in oligotrophic surface oceans, where dominant microbial taxa use light to enhance the acquisition of essential organic substrates(51). However, in the twilight zone, phytoplankton may require a minimum amount of photons for the potential assimilation of exogenous DOC. This process is presumably regulated by cryptochrome-mediated signal cascades(52) and metabolic activity, as indicated by the upregulation of cryptochrome genes, transporters and solute carriers in transcriptomic data and the observed downward trend in DOC concentrations (Fig. 3 and 4). No significant difference in DOC assimilation between high-pressure blue light and dark conditions, likely due to limited culture duration, renders this inference tentative.

### Broader ecological significance

Diatoms are conventionally considered surface-originated particles with negligible activity below the euphotic zone. Accordingly, the abundant deep-water *Chaetoceros* is typically attributed to passive vertical export. This genus can form efficient sinking vectors via resting spores, and its short-lived, boom-and-bust surface blooms create temporal decoupling between surface production and deep-water accumulation(53). Our results refine this view: *Chaetoceros* spp. (represented by DS1) are widely distributed across the twilight zone, with deep-water abundances often comparable to or higher than surface populations (Fig. 5). Integrative multi-omics and physiological evidence collectively confirm that deep-sea *Chaetoceros* cells are metabolically active rather than physiologically inactive detritus. While passive sinking and temporal decoupling cannot be fully ruled out, active twilight-zone inhabitation provides a plausible alternative explanation for their unique vertical distribution. The widespread expression of PF00009, cytochrome and rhodopsin genes of *Chaetoceros* indicates sustained cellular functioning, further ruling out the possibility that these deep populations predominantly consist of transcriptionally silent resting stages. Combined with previous records of *Chaetoceros* enrichment in sinking fluxes and deep-sea sediments, as well as frequent observations of intact deep-water diatom cells, our data support the persistent occurrence of this genus in the deep ocean (12, 53, 54). Nonetheless, we acknowledge inherent limitations in interpreting *in-situ* metabolic status solely from omics datasets. Deep-sea transcripts may partially derive from recently exported surface cells, and cDNA:DNA ratios only reflect relative rather than absolute metabolic activity(55). However, the rapid light-induced physiological reactivation of DS1 demonstrates that mesopelagic *Chaetoceros* cells retain physiological tolerance to prolonged darkness and high hydrostatic pressure. Such stress resilience enables them to actively persist in the twilight zone, instead of merely existing as inert, dormant sinking particles. Collectively, these results support the presence of a metabolically viable subpopulation of *Chaetoceros* in the twilight zone, while we emphasize that the present evidence does not corroborate widespread and continuous *in situ* growth of deep-sea diatoms across mesopelagic habitats.

Several diatom taxa possess adaptive traits highly analogous to those observed in strain DS1, including prolonged dark survival capacity(29, 31, 56), osmotrophic growth under dim light(57, 58), and positive physiological responses to blue light(59). Such adaptations may enable certain cell subpopulations to maintain basal metabolism or to rapidly resume metabolic activity when favorable conditions are restored, whereas other cells may enter a low-metabolism or dormant state. Therefore, diatom populations in the twilight zone may be physiologically heterogeneous, encompassing recent export, low-metabolism persistence, intermittent activity and dormancy, highlighting a continuous physiological spectrum ranging from hypometabolic persistence to swift functional reactivation.

These findings extend the biological-pump paradigm by demonstrating that particulate export may be accompanied by the persistence and intermittent reactivation of viable phytoplankton. Cells retained in the twilight zone may contribute to carbon transformation and nutrient cycling, while mesoscale eddies and wind-driven upwelling may facilitate the return of viable cells to the euphotic zone. Such bidirectional biological transport and physiological heterogeneity are likely underrepresented in oceanographic models(53) that treat the twilight zone primarily as a conduit for particulate carbon export. DS1 therefore provides a mechanistic example of how specialized adaptations may sustain eukaryotic plankton below the euphotic zone and contribute to the diversity and biogeochemical functioning of microbial communities in the global ocean(60, 61). More broadly, these findings underscore the need to resolve the physiological states and ecological functions of phytoplankton inhabiting the ocean twilight zone.

## Materials and Methods

### Strain Isolation and Cultivation

*Chaetoceros* sp. DS1 was isolated from seawater collected at 1000 m depth in the Western Pacific Ocean(28). We verified that DS1 was not derived from upper-water-layer contamination through rigorous sampling protocols, contamination control measures, and the well-documented occurrence of *Chaetoceros* in deep-sea environments(28). Algae isolated from surface seawater were used as controls, including *Phaeodactylum tricornutum* CCMP2561 (originating from the North Atlantic Ocean, obtained from the Provasoli-Guillard National Center for Marine Algae and Microbiota, NCMA), *Nannochloropsis* sp. JH01 (isolated from the coastal seawater, China), and *Chaetoceros muelleri* CCMA318 (isolated from the coastal surface seawater, China). Axenicity of these algae was confirmed by 16S rRNA gene PCR (primers 27F/1492R) and microscopic observation for bacterial contamination over a 1-month culture period. Axenic algae were maintained in autoclaved F/2 medium at 22°C under a 12:12 h light: dark cycle at 40 μmol photons m^-2^ s^-1^. Irradiance was measured with a HopooColor Technology Co., Ltd OHSP-350P spectroradiometer. The exponential-phase cells were used for the following experiments.

### Experimental Setups for Simulated Deep-Sea Conditions

All experiments were conducted in biological triplicate with an overview of treatments provided in Fig. S11.

### Experiment 1: Response to simulated water column layers

To assess DS1’s responses across oceanic depth layers, we simulated conditions for the euphotic layer (surface), 500 m, and 1000 m depths using in situ profiles of temperature, light, and hydrostatic pressure(18): (1) Euphotic layer (Surf-Normal): white light (40 μmol photons m^-2^ s^-1^) + 0.1 MPa (atmospheric pressure) + 22°C. (2) 500 m depth (500m-HP-Dark): Darkness + 5 MPa (generated via a high-pressure vessel; Nantong Feiyu Petroleum Technology Development Co., Ltd., China) + 12°C. (3) 1000 depth (1000m-HP-Dark): Darkness + 10 MPa + 5°C. DS1 and control algal species from the surface ocean were cultured under each condition for 5 days followed by a 3-day recovery under Surf-Normal conditions. A 5-day duration was selected, as the control algae were observed to exhibit decline in cell abundance within this period under darkness and high pressure. Samples were collected at 0 h (initial) and 5 days (treatment endpoint), and post-recovery (3^th^ day) for physiological and transcriptome analyses.

### Experiment 2: Survival and heterotrophic capacity under long-term darkness

To test long-term survival and heterotrophic capacity under prolonged dark conditions (without pressure stress), DS1 cells were cultured in Aquil medium(62) (modified to exclude organic carbon for the “DOC-excluded” treatment) with or without dissolved organic carbon (DOC): (1) Dark-DOC-exc: Darkness + 5°C (DOC excluded); (2) Dark-DOC-add: Darkness + 5°C + 100 μM DOC. DOC was extracted from coastal seawater (Qingdao, China) using C_18_ solid-phase extraction(63), and added at 100 μM (∼2× natural deep-sea DOC levels). Cultures were incubated in complete darkness (wrapped in aluminum foil) for 30 days. Samples were collected at 0, 1, 5, 10, 20, and 30 days for cell counting and physiological assays; transcriptomes were sequenced at 0, 1, 10, and 30 days.

### Experiment 3: Dim blue-light utilization under condition mimicking 500 m depth

To test utilization of residual light in the twilight zone, DS1 was incubated under (1) HP-Blue: dim blue light (450 ± 20 nm, 0.19 ± 0.02 μmol photons m^-2^ s^-1^, Fig. S12) + 5 MPa + 12°C (mimicking light/pressure/temperature at 500 m depth); (2) HP-Dark: darkness + 5 MPa + 12°C (control for aphotic conditions). Cultures were incubated for 5 days, with sampling at 0 and 5 days for physiological and transcriptomic analysis. The 5-day duration was selected to correspond with experiment 1.

To eliminate potential impacts of the duration from pressure release to sample collection, transcriptome samples were collected within 3 min of pressure release, followed by immediate centrifugation for 5 min under complete dark conditions; the resulting pellets were snap-frozen and stored at -80°C right after centrifugation. For Fv/Fm measurement, sampling and subsequent determination were completed within 2-3 min upon collection. For cell abundance analysis, glutaraldehyde was added immediately after sampling for dark fixation. For DOC and nutrient analyses, samples were filtered within 5 min after collection and then stored at -20°C for subsequent assay. To avoid light exposure during subculture sampling and subsample processing, all manipulations were conducted in the dedicated dark room. Notably, *Chaetoceros* sp. DS1 was cultured at a relatively high cell density, not to mimic natural in-situ abundance, but solely to meet the minimum sample requirements for subsequent microscale analyses and ensure detectable, reliable data. This cell density was strictly consistent across all experimental groups, and the potential influence of high abundance on comparing relative changes in physiological and molecular indicators among treatments is therefore expected to be minimal.

### RNA Sequencing and Transcriptomic Analysis

Total RNA was extracted using the mirVana miRNA Isolation Kit (Ambion, USA). The cDNA libraries were constructed using TruSeq Stranded mRNA LTSample Prep Kit (Illumina, USA) and sequenced on the Illumina HiSeqTM 2500 platform (OE Biotech Co., Ltd. China) to generate 150 bp paired-end reads. Raw data were quality-cleaned using Trimmomatic(64) and de novo assembled into transcripts using Trinity v.2.4(65) with default settings. The paired-end clean reads were aligned to our annotated *Chaetoceros muelleri* reference genome (ASM1969354v1) using HISAT2. 2.4(66). FPKM (fragment per kilobase of transcript per million mapped reads) and read counts were calculated using bowtie2(67) and eXpress(68). Differentially expressed genes (DEGs) were identified using the DESeq with FDR-adjusted *P* value < 0.05 thresholds. Kyoto Encyclopedia of Genes and Genomes (KEGG) pathway enrichment analysis of DEGs were conducted using a hypergeometric test (*P* < 0.05).

### Cell Abundance and Morphological Observation

For cell counting, 1.8 mL subsamples were fixed with glutaraldehyde (0.5% v/v final concentration) and stored at 4°C. Cell density was determined in triplicate using a Neubauer hemocytometer under a Leica DM2000 LED microscope (Leica Microsystems, Germany). Morphology was examined via differential interference contrast microscopy, with images captured using a Leica DFC450 C digital camera.

### Photosynthetic Efficiency Measurements

Photosystem II efficiency was measured using a pulse-amplitude modulated chlorophyll fluorometer (Water-PAM, Walz, Germany). For each sample, 3 mL of cell suspension was dark-adapted for 10 min (except dark-maintained samples, measured immediately). Minimum fluorescence (F0) was recorded under low measuring light, followed by a pulse of saturating light (10000 μmol photons m^-2^ s^-1^ for 300 ms) to obtain the maximum fluorescence (Fm). The photochemical efficiency (Fv/Fm) was calculated as Fv/Fm=(Fm-F0)/Fm(69).

### Pigment, Fatty Acid, Organic Matter, and Nutrient Analysis

#### Pigments

10 mL of cell suspension was filtered onto GF/F membranes (25 mm, 0.7 μm, Whatman), rapidly frozen in liquid nitrogen, and stored at -80°C. Pigments were extracted in methanol and analyzed using reversed-phase HPLC(70, 71). Standards (DHI LabProducts, Denmark) were used to quantify chlorophyll a (*Chl a*), fucoxanthin (Fuco), β-carotene (β-Car), diadinoxanthin, and diatoxanthin. The de-epoxidation state (Des) of xanthophyll cycle pigments was calculated as DES = diatoxanthin/(diatoxanthin+ diadinoxanthin)(72).

#### Fatty acid

200 mL of culture was centrifuged (at 8000 rpm for 10 min), lyophilized, and lipids extracted (chloroform:methanol 2:1, v/v). Fatty acid methyl esters (FAMEs) were generated via acid-catalyzed transesterification (5% HCl in methanol, 85°C, 1 h). Heptadecanoic acid (100 μL, 13.6 mg mL^-1^) was added as internal standard. FAMEs were extracted with 1 mL of n-hexane at room temperature for 1 h, and analyzed by GC-MS (Shimadzu GC-MS-TQ8040NX).

#### Cellular organic carbon and nitrogen

20 mL samples were filtered onto pre-combusted (500°C, 5 h) GF/F membranes (25 mm, 0.7 μm, Whatman), exposed to HCl fumes for 24 h to remove inorganic carbon and then dried at 60 °C. Organic carbon and nitrogen were quantified with an EA2000 automatic elemental analyzer (Elementar Analysensysteme Co., Germany).

#### DOC and inorganic nutrients

Filtrates were stored at -20°C. DOC was measured with a total organic carbon analyzer (TOC-L CPH, Shimadzu, Japan). Nitrate (NO_3_-N), and phosphate (PO_4_-P) were measured by spectrophotometry using an Auto-Analyzer (BRAN and LUEBBE AA3, Germany).

### Measurement of Glutathione S-Transferase Activity

Cells were harvested by centrifugation at 8, 000×g for 15 min, washed with deionized distilled water, and then centrifuged again. The pellets were resuspended in 1 mL Tris-HCl buffer (50 mM, pH 7.5) and disrupted using an ultrasonic homogenizer. To prevent overheating during sonication, the tubes were kept on ice throughout the procedure. The homogenate was centrifuged at 4,000×g for 10 min at 4°C, and the supernatant was considered as a cell-free enzyme extract. Protein concentration was determined according to the method described by Bradford(73), with bovine serum albumin as the standard. GST activity was assayed spectrophotometrically at 412 nm by monitoring the conjugation of reduced glutathione with 1-chloro-2,4-dinitrobenzene, according to the method described by Grant et al.(74) with minor modifications. GST activities are reported as U per mg protein (1 U=1 µmol min^-1^). Protein quantification and GST activity were measured with assay kits purchased from Nanjing Jian Cheng Bioengineering Institute (China).

### Global distribution of *Chaetoceros*, Cryptochrome and Rhodopsin Transcripts

To contextualize the ecological relevance of DS1 and its dim blue-light adaptation, we analyzed global distributions of *Chaetoceros* and cryptochrome transcripts using publicly available meta-omics datasets from the Tara Oceans expedition. We accessed Tara Oceans metatranscriptomic data (MATOUv1-T) and taxonomic profiles, which include samples from surface, DCM, and twilight zones (200-1000 m) across global ocean basins. These datasets cover multiple size fractions, with a focus on the 0.2-3 μm fraction (representing ultraphytoplankton) and include annotated eukaryotic transcripts. Raw data and processed taxonomic tables (miTAGs) were retrieved from the European Nucleotide Archive under project accessions corresponding to the MATOUv1-T release(37, 42). From the Tara Oceans miTAG taxonomic tables (derived from 0.2-3 μm size fractions), we first extracted eukaryotic algal taxa. We then retained only stations with paired samples from surface, DCM, and twilight zones, ensuring at least one twilight zone sample per station contained detectable *Chaetoceros* reads. To account for variations in sequencing depth across samples, we normalized *Chaetoceros* abundance within each sample by calculating its relative proportion (%) of total eukaryotic algal reads (i.e., *Chaetoceros* reads/total algal reads × 100). This approach standardized abundance estimates for cross-layer and cross-station comparisons. From the MATOUv1-T metatranscriptomic data, we identified cryptochrome-encoding transcripts by BLASTp alignment (E-value < 1e-10) against *Chaetoceros* sp. DS1’s cryptochrome and rhodopsin protein sequences (Tables S5 and S6)(75). Relative abundance of these transcripts was calculated as a percentage of total eukaryotic transcripts per sample to assess their expression patterns across depth layers. Distribution patterns of *Chaetoceros* and cryptochrome/rhodopsin transcripts across surface, DCM, and twilight zone were visualized using Ocean Data View Software (ODV, https://odv.awi.de/) and depth-specific plots, highlighting stations where *Chaetoceros* was detected in twilight zones.

### Analysis of Tara Oceans metatranscriptomic data by searching PF00009 (i.e., GTP-binding elongation factor family, EF-Tu/EF-1A subfamily)

To address concerns about *Chaetoceros* being passively exported as resting stages, we analyzed MATOUv1-T metatranscriptomic data from the Tara Oceans stations with samples collected cross surface (0.8-5 μm size fraction), deep chlorophyll maximum (DCM, 0.8-5 μm size fraction), and the twilight zone (0.8-3 μm size fraction), using PF00009 as a marker. This Pfam family is among the most highly expressed diatom-associated families in the global ocean, supporting protein synthesis. Selection of different size-fractionated samples was due to the 0.8-5 μm fraction not being sampled in the twilight zone, whereas no samples or annotated data for the 0.8-3 μm fraction were available for the surface and DCM layers. We identified PF00009 (https://www.ebi.ac.uk/interpro/entry/pfam/PF00009/) via BLASTp alignment with an E-value cutoff of < 1e-10 against MATOUv1-T metatranscriptomic data. We further quantified the relative abundance of taxa corresponding to PF00009 at each sampling station, and characterized the community structure of eukaryotic phytoplankton across different sampling stations and water layers based on Kraken analysis using Ocean Gene Atlas v2.0 (https://tara-oceans.mio.osupytheas.fr/ocean-gene-atlas/)(75).

### Statistical Analysis

Data were presented as mean ± standard deviation (SD) of triplicate measurements. Levene’s test was used to verify homogeneity of variances. Significant differences between treatments were assessed via Welch’s t-test (two-tailed) at *p* < 0.05. All statistical analysis was performed in SPSS Statistics.

## Acknowledgments

We thank all members aboard the R/V “Dongfanghong 3” and the team members of the HOV “Jiaolong” manned submersible for their efforts in the sample collection. We also acknowledge the open research cruises NORC2020-06 and NORC2023-801, which were supported by the Ship Time Sharing Project of the National Natural Science Foundation of China (NSFC).

## Data, Code, and Materials Availability

All data and code needed to evaluate and reproduce the results in the paper are present in the paper and/or the Supplementary Materials. There are no new materials generated in the study. The raw transcriptome sequence data reported in this paper have been deposited in the China National Center for Bioinformation (CNCB) GSA database under accession (PRJCA046893).

## Table of contents for supplementary materials

This PDF file includes:

Supplementary Text

Figs. S1 to S12

Other Supplementary Materials for this manuscript include the following:

Tables S1 to S6

