## Supplementary material for "Diatoms maintain metabolic activity in deep ocean twilight zones": 20260728 Supplementary Information: 20260728 Supplementary Information.docx

**This PDF file includes:**

Supplementary Text

Figs. S1 to S12

**Other Supplementary Materials for this manuscript include the following:**

Tables S1 to S6

**Supplementary Text**

**In-situ 18S rRNA transcript sequencing from Northwestern Pacific stations**

Sampling was conducted during a research cruise to the Northwestern Pacific Ocean in October 2025. Vertical water column samples were collected at stations D1 (30°24’N; 145°13’E) and D6 (41°N; 151°13’E), covering the surface layer, deep chlorophyll maximum (DCM), twilight zone, and deep-sea layers. For each depth, 1 L of seawater was sequentially filtered through a 200-μm mesh sieve to remove large particles, followed by filtration onto a 0.2-μm Millipore membrane filter. The collected filters were immediately immersed in RNAlater, flash-frozen in liquid nitrogen, and stored at -80 °C until further processing.

Upon return to the laboratory, total RNA was extracted from the filters. The V4 region of the Eukaryotic 18S rRNA gene (528F: GCGGTAATTCCAGCTCCAA; 706R: AATCCRAGAATTTCACCTCT) was amplified from RNA samples. Amplicon libraries were pooled in equimolar amounts and sequenced on an Illumina MiSeq platform (2 x 300 bp paired-end) at Magigene Biotech Co., Ltd. Raw sequence reads were processed using the package DADA2 in R following the default settings. Here, the reads were filtered with the following parameters: maxN = 0, maxEE = c (5, 5), truncQ = 2, minBoot = 80. Taxonomic assignment of ASVs of 18S rRNA genes was performed using the Silva 138.2 database. Non-phytoplankton taxa were manually excluded from the resulting taxonomic table, and the remaining reads were rarefied to generate a normalized relative abundance table for eukaryotic phytoplankton. Focused analyses were performed to visualize the relative proportion of diatoms within total phytoplankton and the relative contribution of *Chaetoceros* within the diatom community across depth layers. Photosynthetically active radiation (PAR) data were obtained from concurrent CTD measurements.

**Calculation of light attenuation by syringes in high-pressure algal cultivation experiments**

Algal cells were cultivated in syringes placed inside a **high-pressure cultivation apparatus**. To accurately determine the actual light intensity received by the algae, it was essential to correct for the light attenuation caused by the syringe walls. Since direct measurement of light intensity inside the 10 mL syringes used in the experiments was not feasible, we first quantified the light attenuation in 100 mL syringes made of the identical plastic material. Based on the Beer-Lambert law, we assumed that light absorption by the syringe material is proportional to its wall thickness. Using this assumption, we then calculated the theoretical light attenuation for the 10 mL syringes. The detailed calculation procedure is summarized below:

#### Step 1: Average Light Intensity Measurement for 100 mL Syringes

- **Blank control (no syringe):** replicate measurements yielded an average light intensity of I_blank_ = 0.15375.
- **With 100 mL syringe:** replicate measurements yielded an average light intensity of I_with_ = 0.098.

#### Step 2: Calculation of 100 mL Syringe Optical Parameters

1. **Measured Transmittance (T_100mL_):**

T_100mL_ = I_blank_/I_with_ = 0.6374

2. **Measured Attenuation Rate (A_100mL_):**

A_100mL_=1−T_100mL_=36.26%

3. **Material Absorption Coefficient (α):**

Using the wall thickness of 1.0 mm for the 100 mL syringe, the intrinsic absorption coefficient of the plastic material was calculated as α = 0.4504 /mm. This value is universal for both 10 mL and 100 mL syringes of the same batch.

#### Step 3: Theoretical Calculation for 10 mL Syringes

Using the universal absorption coefficient α and the standard wall thickness of 0.9 mm for 10 mL syringes:

1. **Theoretical Transmittance (T_10mL_):**

T_10mL_=e^-α d^ = 0.6667

2. **Theoretical Attenuation Rate (A_10mL_):**

A_10mL_=1−T_10mL_=33.33%

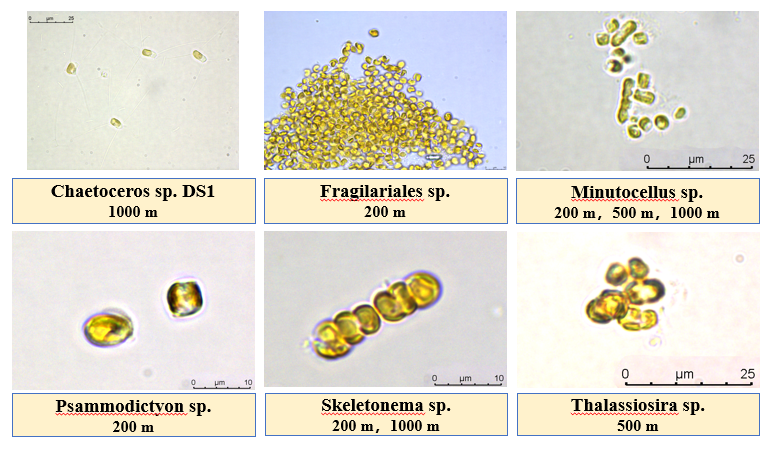

Figure S1. Morphology and taxonomic identification of diatom strains isolated from the Northwest or West Pacific Ocean.

Scale bars are indicated in the figure. Isolation depths are specified below each taxon label.

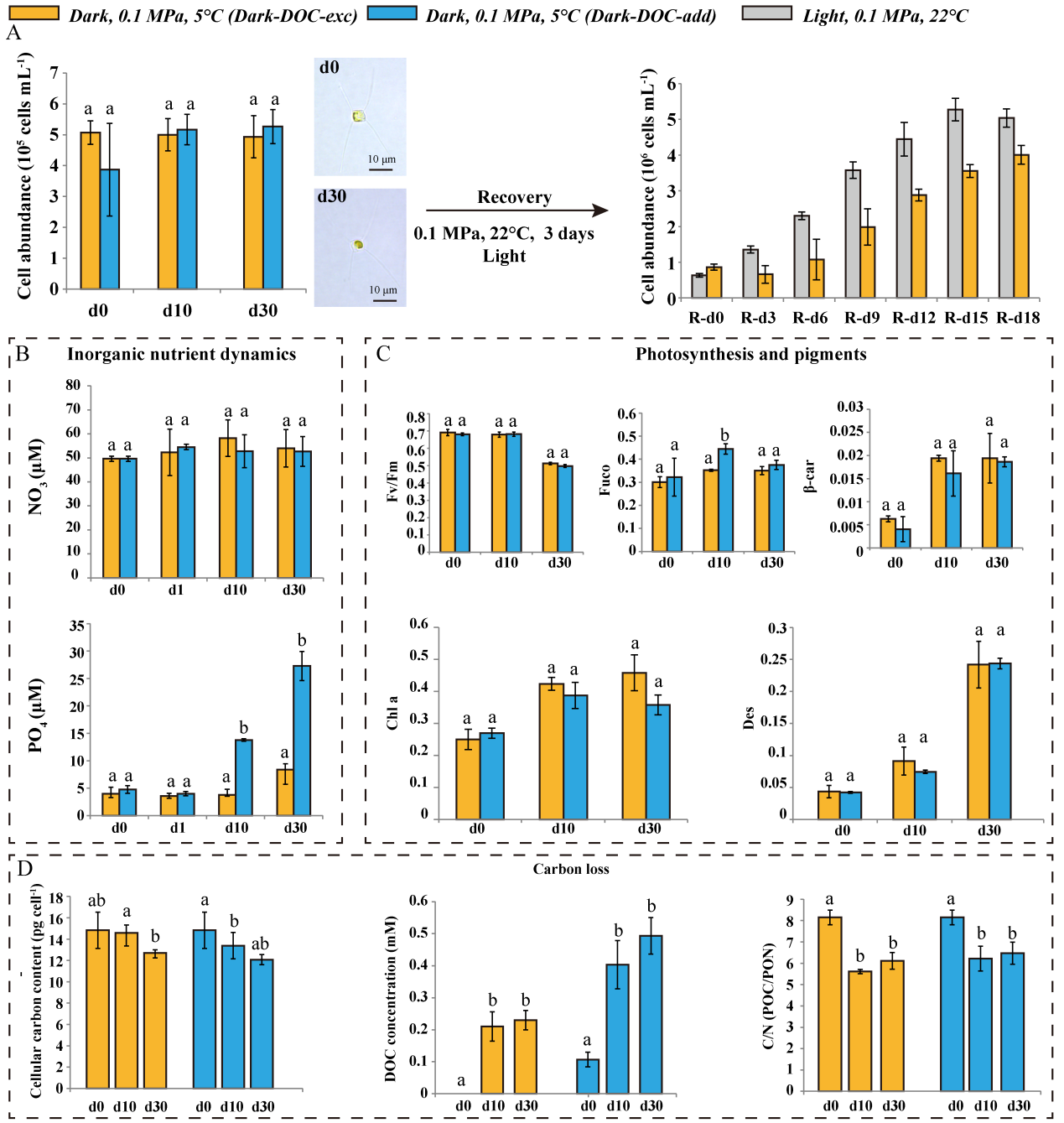

Figure S2. Physiological and biochemical characteristics of Chaetoceros sp. DS1 under long-term darkness with or without exogenous dissolved organic carbon (DOC)

Color legend: Orange refers to Dark-DOC-exc (darkness, 0.1 MPa, 5°C, no exogenous DOC), blue refers to Dark-DOC-add (darkness, 0.1 MPa, 5°C, exogenous DOC), and gray refers to Light (white light, 0.1 MPa, 22°C, reference condition). (A) Left panel shows cell abundance dynamics under Dark-DOC-exc and Dark-DOC-add treatments over 30 days, and stable cell counts were maintained without formation of dormant spores (microscopic insets, day0 (d0) and Day30 (d30); scale bar: 10 μm). Right panel presents recovery of cell abundance during an 18-day re-illumination period under ambient light and pressure (0.1 MPa, 22°C). (B) Temporal changes in nitrate (NO_3_^-^, top) and phosphate (NO_3_^-^, bottom) concentrations (μM) across 30 days. (C) Variations in photosynthetic efficiency (Fv/Fm), total pigment content, chlorophyll a (Chl a), fucoxanthin (Fuco), β-carotene, and de-epoxidation state (Des) levels over 30 days. (D) Changes in cellular carbon content (fg cell^-1^), DOC concentration in culture media (μM), and C/N ratio across 30 days. Letters denote statistical significance (Welch’s t-test, *p* < 0.05).

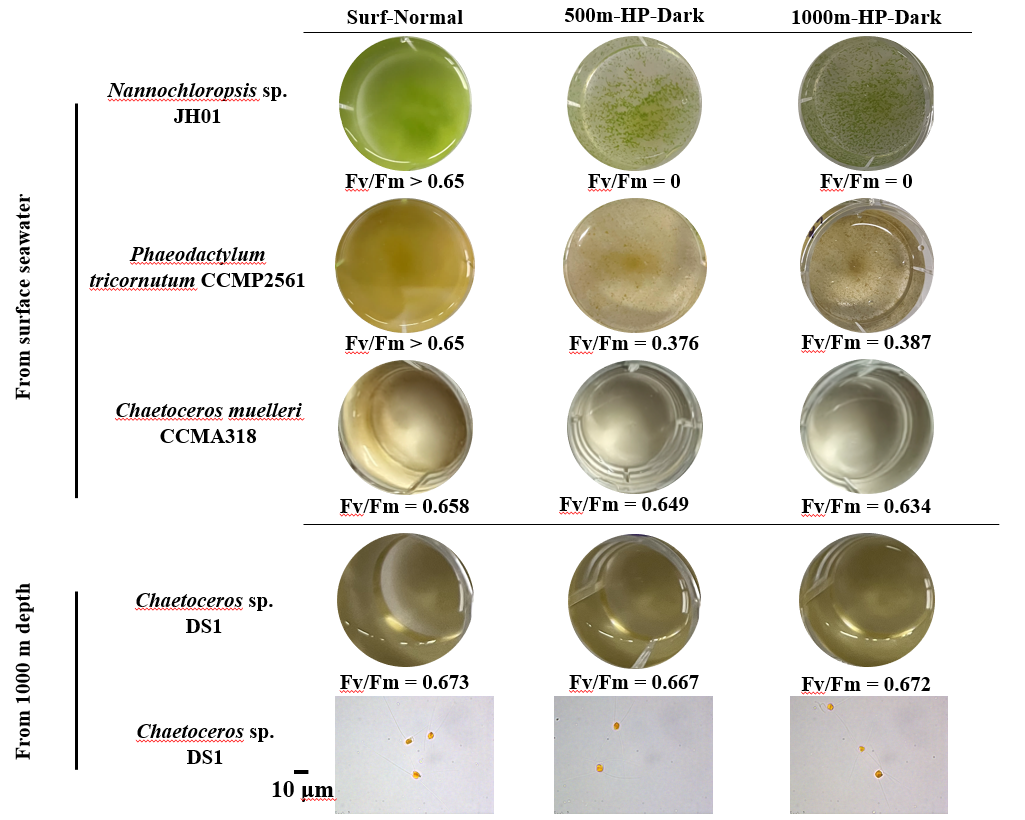

**Figure S3. Growth and photosynthetic efficiency (Fv/Fm) responses of *Chaetoceros* sp*.* DS1 under simulated deep-sea conditions.**

Surf-Normal: white light, 0.1 MPa, 22°C; 500m-HP-Dark: mimicking 500 m depth, darkness, 5 MPa, 12°C; 1000m-HP-Dark: mimicking 1000 m depth, darkness, 10 MPa, 4°C. Three control algal strains collected from the surface seawater are *Nannochloropsis* sp. JH01, *Phaeodactylum tricornutum* CCMP2561 and *Chaetoceros muelleri* CCMA318. The microscopy images of *Chaetoceros* sp. DS1 cells were below, scale bar = 10 μm.

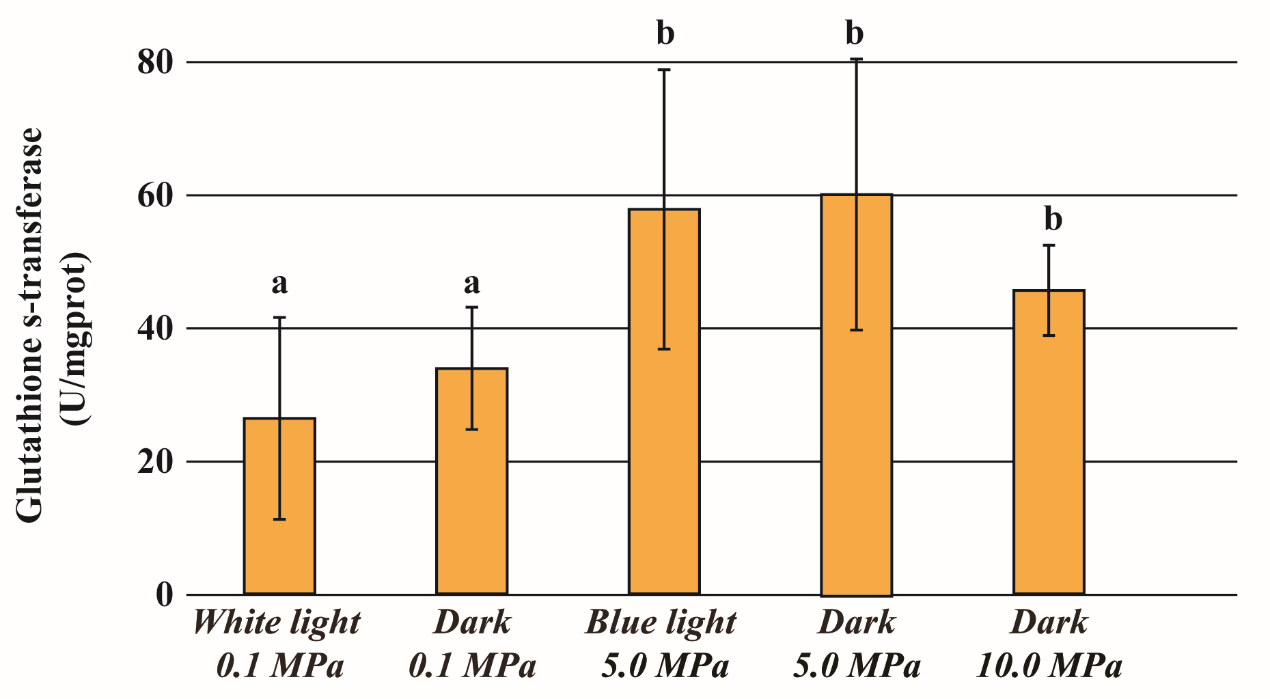

**Figure S4. Glutathione s-transferase activity of *Chaetoceros* sp. DS1 under simulated environmental conditions.**

These conditions include white light + 0.1 MPa, darkness + 0.1 MPa, darkness + 5 MPa, darkness + 10 MPa, and dim blue light (0.19 ± 0.02 μmol photons m^-2^ s^-1^) +5 MPa. Error bars denote standard deviation; letters indicate statistical significance (Welch’s t-test, *p* < 0.05).

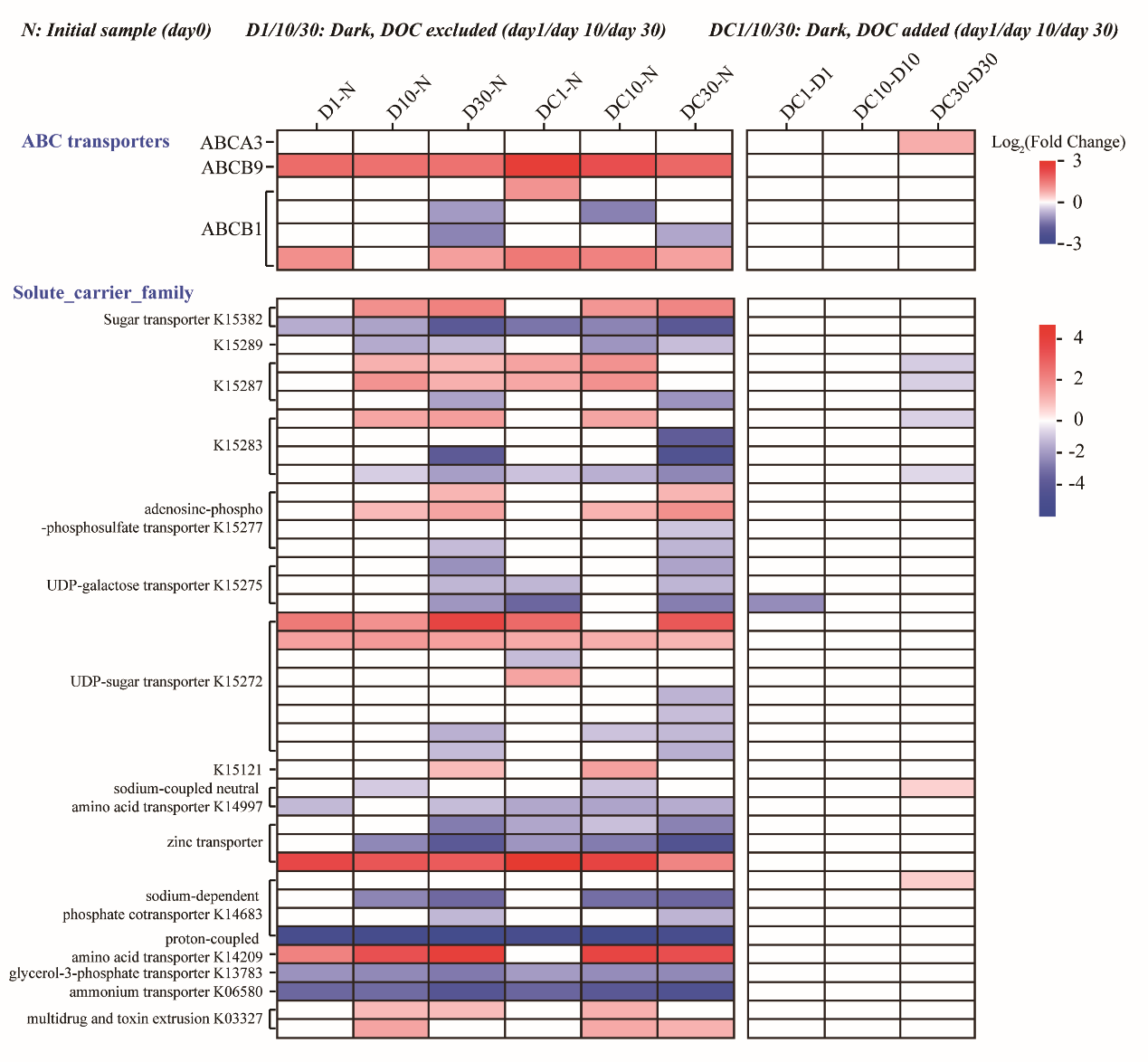

**Figure S5. Transcriptional profiles of transporter-related genes in *Chaetoceros* sp. DS1 under long-term darkness with or without exogenous dissolved organic carbon (DOC)**

Experimental groups: N (initial sample, day 0), D1/10/30 (darkness, DOC excluded, sampled at Day 1, 10, 30), DC1/10/30 (darkness, DOC added, sampled at Day 1, 10, 30). Heatmaps illustrate log_2_ fold change in gene expression to initial (Day 0) levels for genes encoding ABC transporters and solute carrier family members (subtypes include transporters for sugars, phosphate, amino acids, metals, and UDP-sugars). Color gradient: Red denotes upregulation, blue denotes downregulation, with color intensity corresponding to fold change magnitude (scale bars indicate log_2_fold change range).

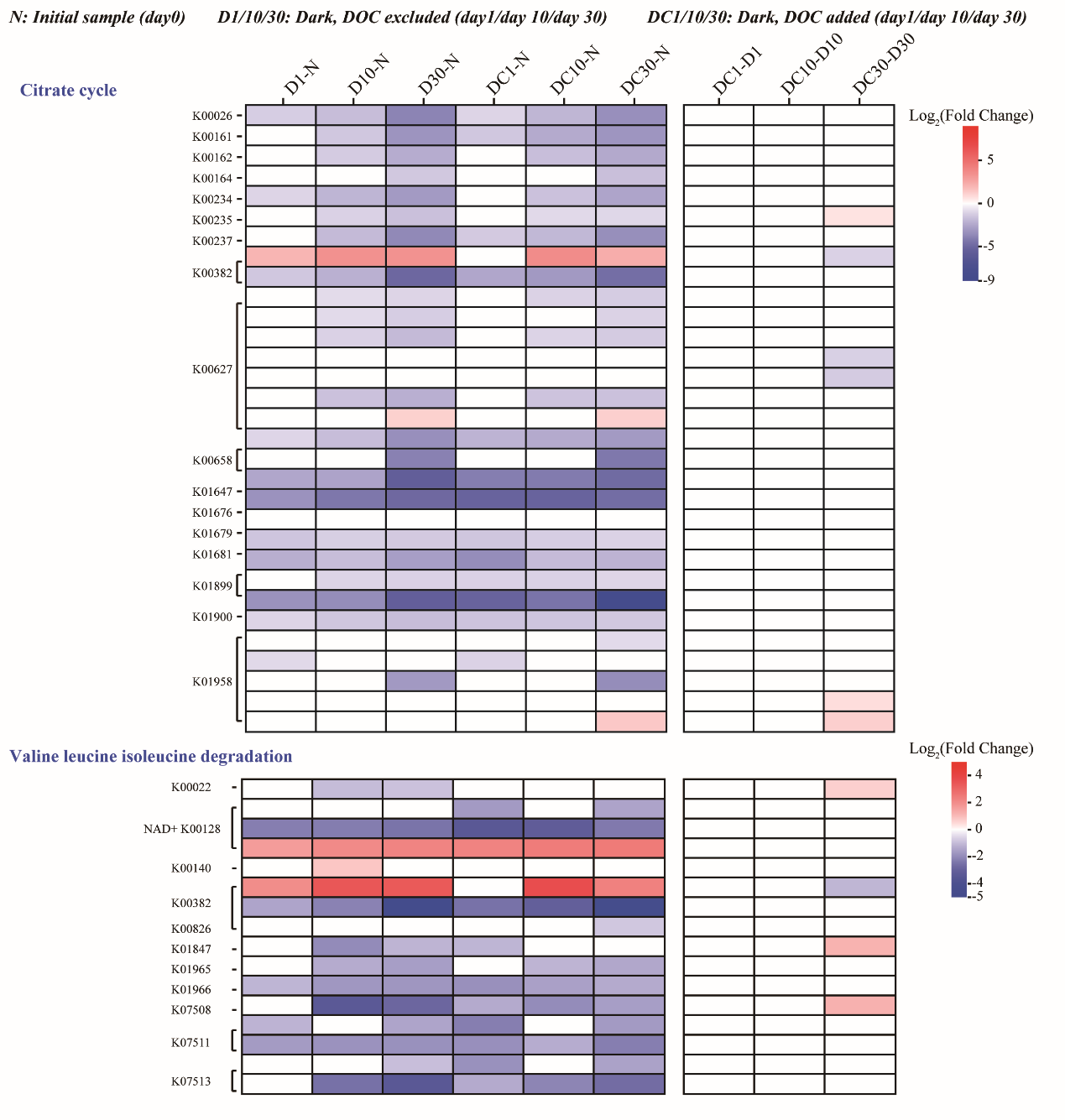

**Figure S6. Transcriptional profiles of tricarboxylic acid cycle and amino acid degradation pathways in *Chaetoceros* sp. DS1 under prolonged darkness with or without exogenous dissolved organic carbon (DOC)**

Experimental groups: N (initial sample, day 0), D1/10/30 (darkness, DOC excluded, sampled at Day 1, 10, 30), DC1/10/30 (darkness, DOC added, sampled at Day 1, 10, 30). Heatmaps illustrate log_2_ fold change in gene expression for the citrate cycle (top) and valine, leucine, and isoleucine degradation (bottom) pathways. Color gradients: Red denotes upregulation, blue denotes downregulation, with color intensity corresponding to fold change magnitude (scale bars indicate log_2_ fold change range for each pathway module).

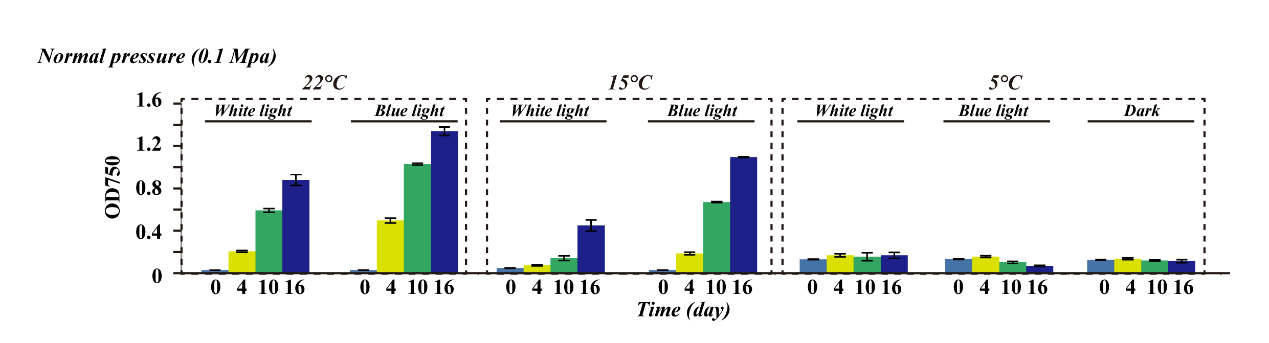

**Figure S7. Physiological performance of *Chaetoceros* sp. DS1 at ambient pressure under different temperatures (22°C, 15°C, 5°C) and light regimes (white light vs blue light).** error bars represent standard deviation.

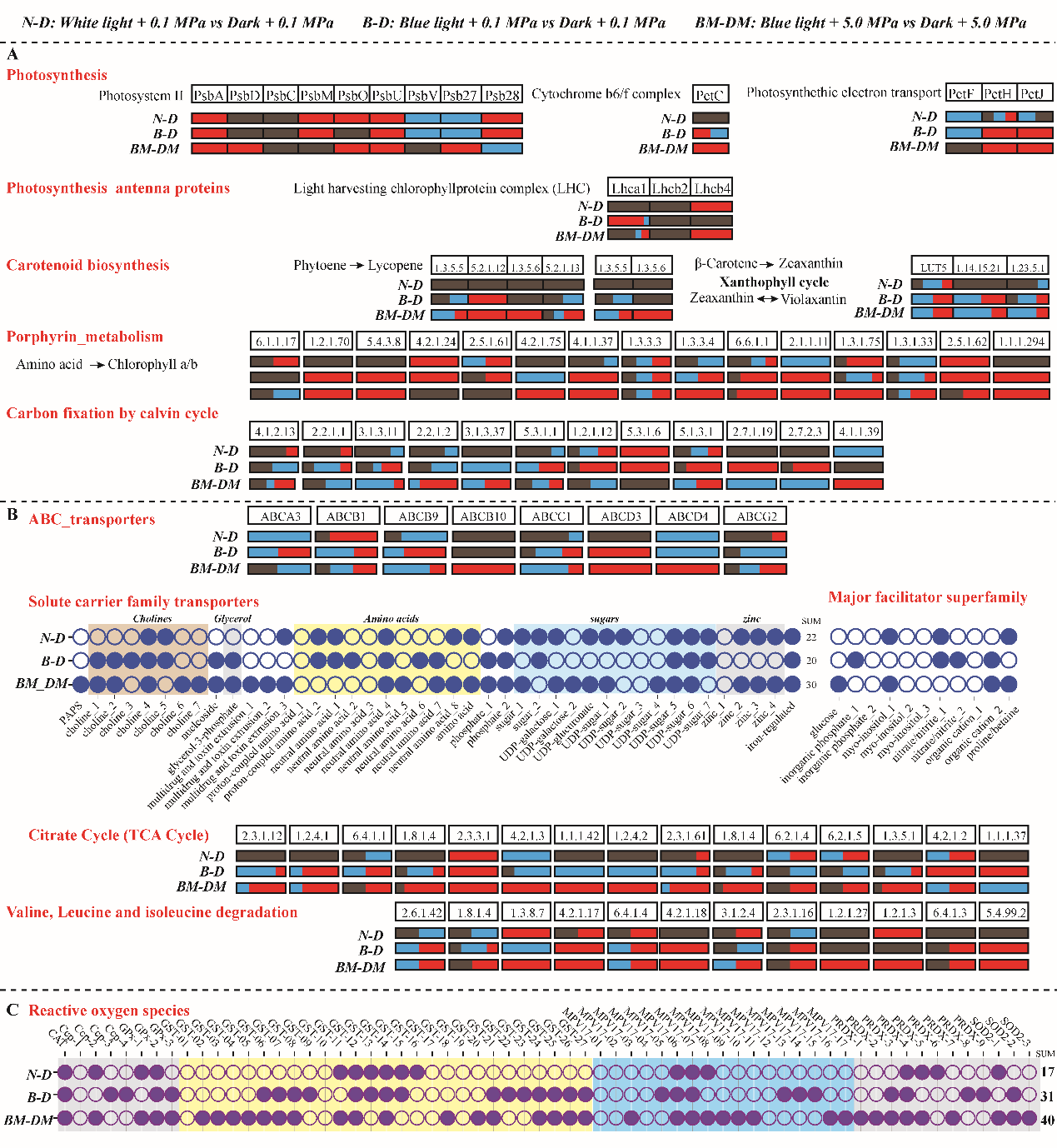

**Figure S8. Transcriptional dynamics of *Chaetoceros* sp. DS1 under dim blue light and pressure treatments**

**Experimental design (top)**: Transcriptomic contrasts include N-D (White light + 0.1 MPa vs darkness + 0.1 MPa), B-D (dim blue light + 0.1 MPa vs darkness + 0.1 MPa), and BM-DM (dim blue light + 5.0 MPa vs darkness + 5.0 MPa). **Visualization scheme:** In the boxes, red indicates significant upregulation, blue indicates significant downregulation, and brown indicates non-significant expression change. Each horizontal bar represents a gene cluster, with segment proportions reflecting the fraction of genes in each category. Circles indicate gene-level responses: solid circle indicates significantly upregulated genes; open circle indicates downregulated or non-significantly changed genes. **Functional categories:** (A) Photosynthesis and pigment metabolism: Photosystem I/II, cytochrome b_6_f complex, photosynthetic electron transport, light-harvesting chlorophyll a/b complexes, carotenoid biosynthesis (phytoene to lycopene, xanthophyll cycle), porphyrin metabolism (amino acid to chlorophyll a/b), and carbon fixation. (B) Transporters and central carbon/nitrogen metabolism: ABC transporters, solute carrier family transporters (amino acids, carbohydrates), major facilitator superfamily, tricarboxylic acid cycle, and valine/leucine/isoleucine degradation. (C) Reactive oxygen species scavenging: Glutathione peroxidase (GPx), peroxiredoxin (PRDX), catalase (CAT), cytochrome c peroxidase (CCP), glutathione S-transferase (GST), mitochondrial inner membrane protein 17 (MPV17), and superoxide dismutase (SOD).

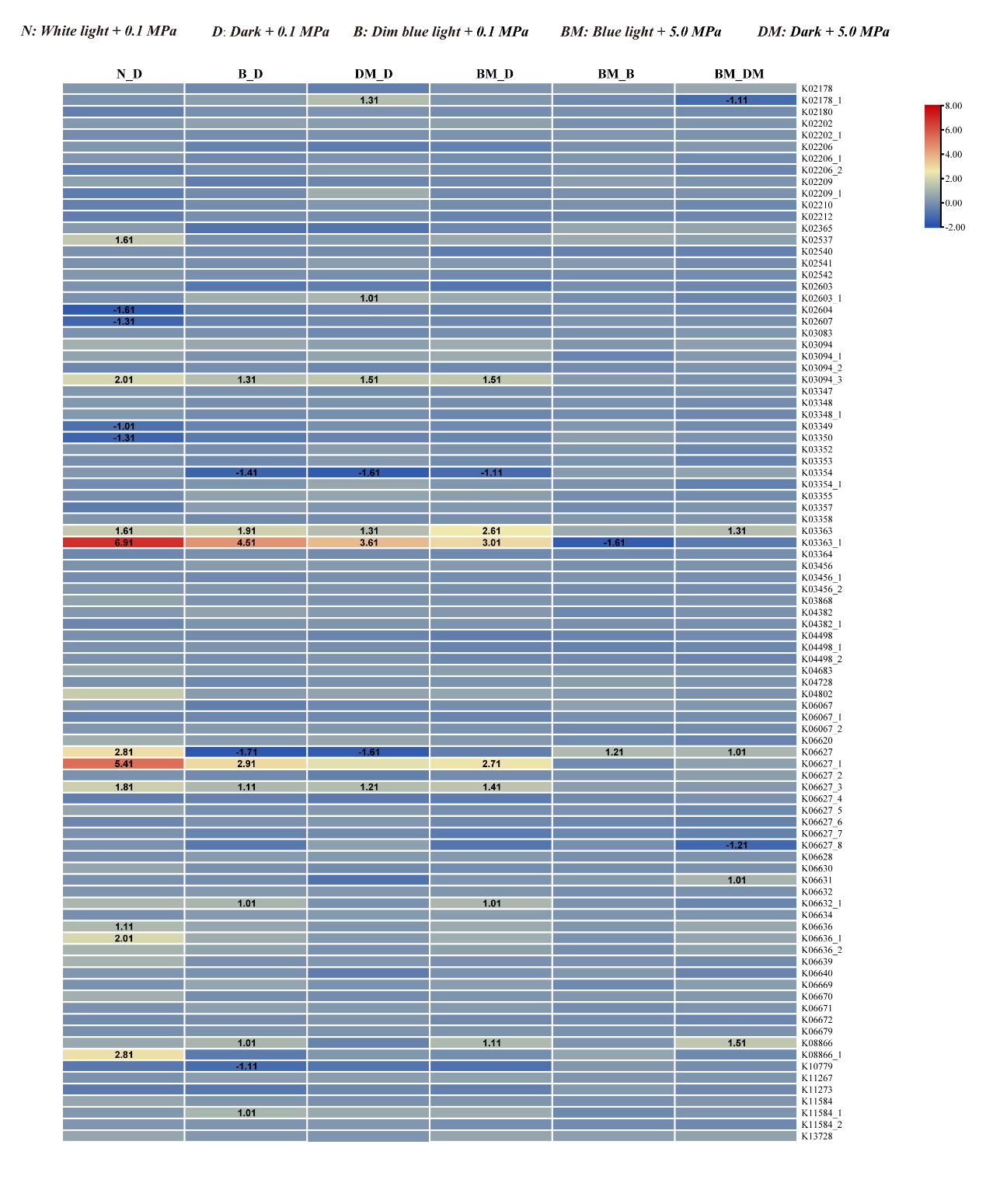

**Figure S9. Transcriptional responses of cell cycle-related genes to different environmental treatments.** N-D (White light + 0.1 MPa vs darkness + 0.1 MPa), B-D (dim blue light + 0.1 MPa vs darkness + 0.1 MPa), DM-D (darkness + 5.0 MPa vs darkness + 0.1 MPa), BM-D (dim blue light + 5.0 MPa vs darkness + 0.1 MPa), BM-B (dim blue light + 5.0 MPa vs dim blue light + 0.1 MPa) and BM-DM (dim blue light + 5.0 MPa vs darkness + 5.0 MPa). Differentially expressed genes were defined as |log_2_FoldChange| > 1 and *p* < 0.05, and corresponding expression response values were marked.

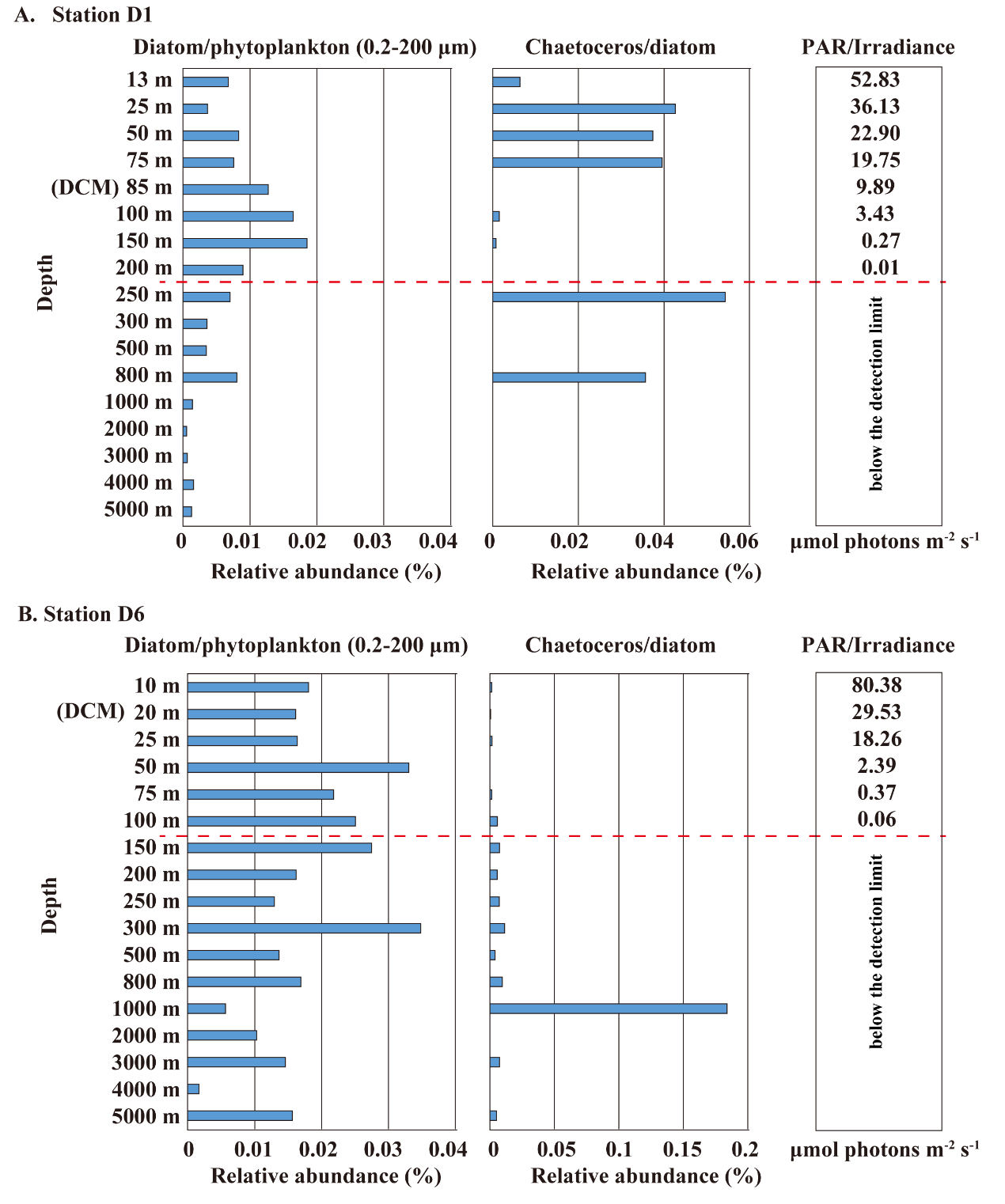

**Figure S10. Vertical distribution of diatom community composition (0.2-200 μm) based on 18S rRNA gene analysis at two Northwestern Pacific stations**

(A) Station D1 and (B) Station D6: vertical profiles showing the relative abundance of diatoms within the total phytoplankton community (left) and the relative contribution of *Chaetoceros* within the diatom community (right). Samples were collected across the surface layer, deep chlorophyll maximum (DCM), twilight zone and deep-sea layers during the Northwestern Pacific research cruise. In-situ light intensity data were obtained from concurrent CTD measurements.

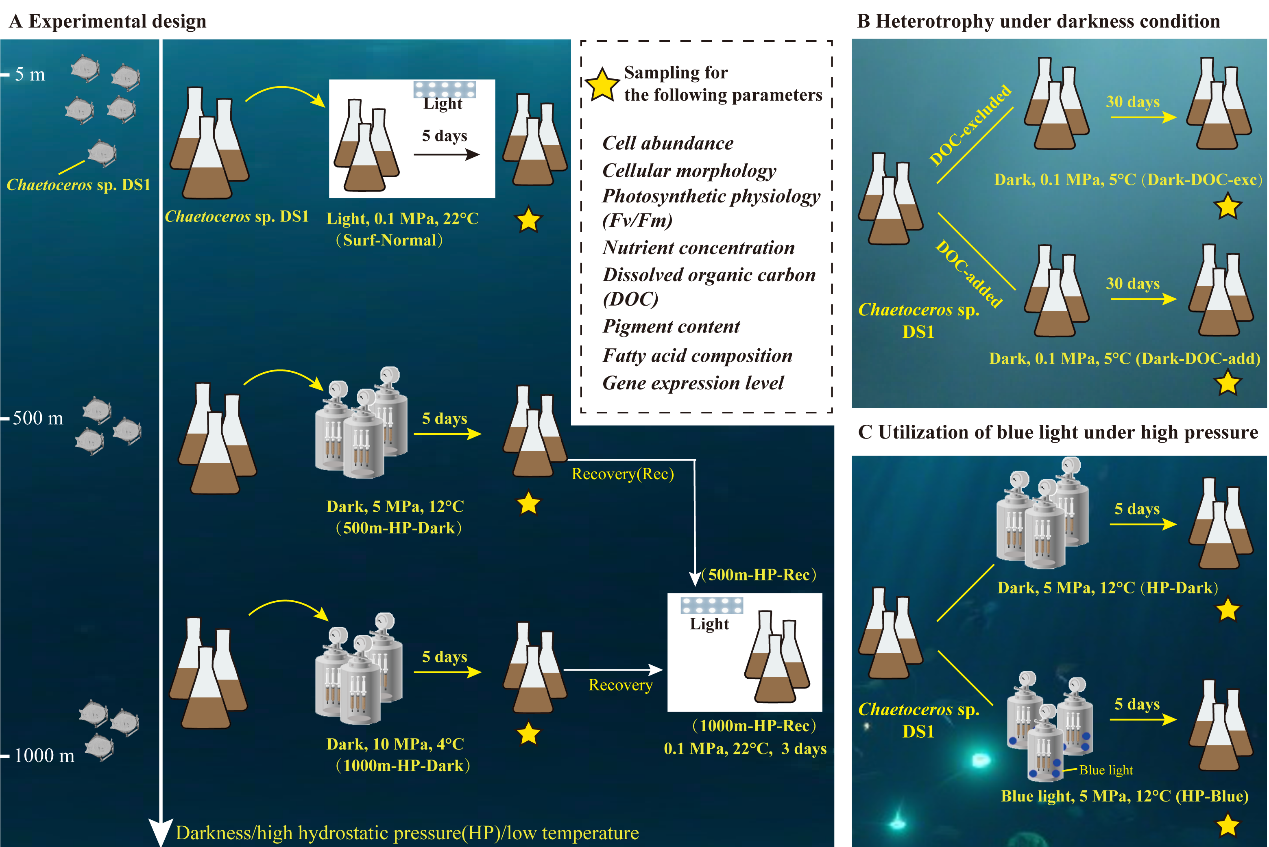

**Figure S11. Experimental design to investigate *Chaetoceros* sp. DS1 responses to twilight zone** **conditions**

(A) **Simulated water column layers**. *Chaetoceros* sp. DS1 was incubated for 5 days under three conditions: Surf-Normal (white light, 0.1 MPa, 22°C), 500m-HP-Dark (darkness, 5 MPa, 12°C), and 1000m-HP-Dark (darkness, 10 MPa, 4°C). A 3-day “Recovery” phase (under ambient light/pressure) tested algal resilience following high-pressure/dark stress. (B) **Survival and heterotrophic capacity under long-term dark conditions**. DS1 was incubated for 30 days in darkness (5°C, 0.1 MPa) with with either no exogenous DOC (Dark-DOC-exc) or 100 μM DOC supplementation (Dark-DOC-add); yellow stars mark sampling timepoints. (C) **Dim blue-light utilization under condition mimicking 500 m depth**. DS1 was exposed for 5 days to HP-Dark (darkness, 5 MPa, 12°C) or HP-Blue (0.19 ± 0.02 μmol photons m^-2^ s^-1^ blue light, 5 MPa, 12°C). Yellow stars indicate sampling points for measuring cell abundance, morphology, photosynthetic physiology (Fv/Fm), nutrient and DOC concentrations, pigment composition, fatty acid profiles, and transcriptomes.

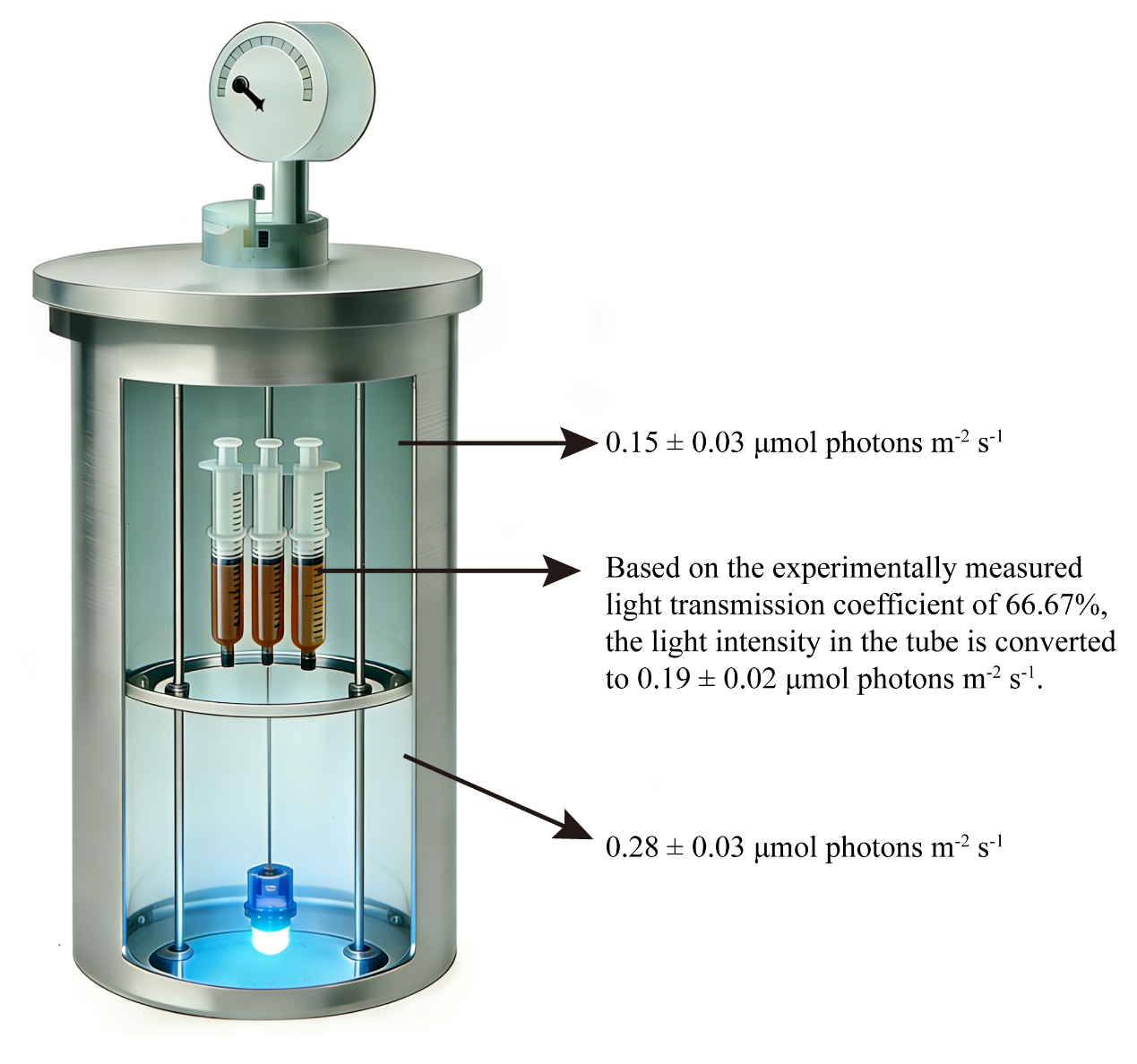

**Figure S12. Experimental setup and light intensity value for diatom incubation.**

Schematic of the custom-built incubation system. The setup consists of a light source at the bottom, and syringes containing diatom cultures suspended within the tube. Light intensity was measured at three key positions. Above the syringe array: 0.15 ± 0.03 photons m^-2^ s^-1^. Within the diatom-containing syringes: 0.19 ± 0.02 μmol photons m^-2^ s^-1^ (calculated from a measured light transmission coefficient of 66.67% through the syringe material). Below the syringe array (near the light source): 0.28 ± 0.03 μmol photons m^-2^ s^-1^.

**Table S1. KEGG enrichment of down-regulated differentially expressed genes under darkness conditions**

**Table S2. KEGG enrichment of up-regulated differentially expressed genes under darkness conditions**

**Table S3 Transcriptional dynamics of genes involved in photosynthetic complexes, pigment biosynthesis and fatty acid metabolism under simulated high-pressure and dark conditions**

**Table S4 The relative abundance of ultraphytoplankton (0.2-3 μm) across surface, Deep Chlorophyll Maximum (DCM), and Twilight zones (Twz; 200-1000m) at Tara Oceans stations**

**Table S5 *Chaetoceros* sp. DS1’s cryptochrome gene and transcript sequences**

**Table S6 *Chaetoceros* sp. DS1’s rhodopsin gene and transcript sequences**
